# RNA recognition by minimal ProQ from *Neisseria meningitidis*

**DOI:** 10.1101/2024.07.24.604975

**Authors:** Maciej Basczok, Mikołaj Olejniczak

**Affiliations:** Institute of Molecular Biology and Biotechnology Faculty of Biology, Adam Mickiewicz University Uniwersytetu Poznańskiego 6, 61-614 Poznań, Poland

**Keywords:** *Neisseria* ProQ, the FinO domain, bacterial regulatory RNA, RNA-binding proteins in bacteria

## Abstract

*Neisseria meningitidis* minimal ProQ is a global RNA binding protein belonging to the family of FinO-domain proteins. The *N. meningitidis* ProQ consists only of the FinO domain accompanied by short N- and C-terminal extensions. To better understand how this minimal FinO-domain protein recognizes RNAs, we compared its binding to seven different natural RNA ligands of this protein. Next, two of these RNAs, *rpmG*-3’ and AniS, were subject to further mutational studies. The data showed that *N. meningitidis* ProQ binds the lower part of the intrinsic transcription terminator hairpin, and that the single-stranded sequences on the 5’ and 3ʹ side of terminator stem are required for tight binding. However, the specific lengths of 5’ and 3ʹ RNA sequences required for optimal binding differed between the two RNAs. Additionally, our data show that the 2ʹ-OH and 3ʹ-OH groups of the 3ʹ terminal ribose contribute to RNA binding by *N. meningitidis* ProQ. In summary, the minimal ProQ protein from *N. meningitidis* has generally similar requirements for RNA binding as the isolated FinO domains of other proteins of this family, but differs from them in detailed RNA features that are optimal for specific RNA recognition.

## INTRODUCTION

FinO-domain proteins are a diverse family of RNA-binding proteins, which are present in many proteobacteria (Glover et al. 2015; Attaiech et al. 2017; Olejniczak and Storz 2017; Holmqvist and Vogel 2018; Holmqvist et al. 2020). Besides the RNA-binding FinO domain these proteins also often contain N-terminal or C-terminal extensions, which contribute to their physiological functions (Arthur et al. 2003; Attaiech et al. 2016; El Mouali et al. 2021b; Rizvanovic et al. 2021). The FinO-domain proteins bind to regulatory RNAs and mRNAs (van Biesen and Frost 1994; Attaiech et al. 2016; Smirnov et al. 2016; Holmqvist et al. 2018; Melamed et al. 2020), and contribute to the regulation of important physiological processes, including the F plasmid transfer (van Biesen and Frost 1992; Glover et al. 2015), natural transformation (Attaiech et al. 2016), adaptation to available nutrients (El Mouali et al. 2021b), motility (Rizvanovic et al. 2021), and bacterial virulence (Westermann et al. 2019; Rizvanovic et al. 2022; Bergman et al. 2024). In bacterial cells FinO-domain proteins coexist with a matchmaker protein Hfq (Updegrove et al. 2016; Kavita et al. 2022; Malecka and Woodson 2024), but ProQ and Hfq mostly recognize different RNA targets (Holmqvist et al. 2018; Melamed et al. 2020). Although all FinO-domain proteins have the same RNA binding domain, there are wide differences between them in RNA recognition, because some bind few RNAs (van Biesen and Frost 1994; Attaiech et al. 2016; Gerovac et al. 2020; El Mouali et al. 2021a), while others are global RNA binders (Holmqvist et al. 2018; Bauriedl et al. 2020; Melamed et al. 2020).

Intrinsic transcription terminators are the binding sites of FinO-domain proteins in small RNAs and mRNAs (Arthur et al. 2011; Attaiech et al. 2016; Holmqvist et al. 2018; Bauriedl et al. 2020; Melamed et al. 2020; Stein et al. 2020; Kim et al. 2022). The FinO domain forms a compact shape with clearly defined convex and concave surfaces (Ghetu et al. 2000; Chaulk et al. 2010; Gonzalez et al. 2017; Immer et al. 2020; Kim et al. 2022). The concave face has been shown as the RNA binding site in F-like plasmid FinO (Ghetu et al. 2002), in *Escherichia coli* ProQ (Pandey et al. 2020; Stein et al. 2023), and in *Legionella pneumophila* RocC proteins (Kim et al. 2022). The crystal structure of the isolated FinO domain of *Legionella pneumophila* RocC with the terminator of RocR sRNA showed how the terminator hairpin with the 3ʹ tail binds to the concave face of the FinO domain (Kim et al. 2022). Particularly important for the interaction are two regions of the protein. One of them is a group of amino acids in α-helix 5, which side chains contact the phosphor-sugar backbone of the lower part of the terminator stem. The other region is a pocket on the concave face where side chains of conserved tyrosine and arginine together with other residues contact two terminal nucleotides of the 3ʹ tail of RocR (Kim et al. 2022).

Besides the FinO domain other regions can also contribute to RNA binding, but such regions are not present in all proteins from this family (Attaiech et al. 2017; Olejniczak and Storz 2017). The N-terminal extension of the F-like plasmid FinO protein contributes to RNA binding and strand exchange (Ghetu et al. 2002; Arthur et al. 2003). Additionally, it was observed that while the isolated FinO domain of *E. coli* ProQ protein can bind only to RNAs containing intrinsic transcription terminators or similar structures on their 3ʹ ends, the full-length ProQ, which contains a positively charged linker, can bind well also to RNAs devoid of such structures (Stein et al. 2020). The role of the ProQ linker in RNA binding was also proposed using the hydrogen-deuterium exchange studies (Gonzalez et al. 2017). These data suggest that N- or C-terminal extensions can contribute to RNA binding by FinO domain proteins. However, not all proteins from this family contain such additional regions, which raises the question whether such minimal proteins consisting only of FinO domains recognize RNAs in the same way as the FinO domains of proteins containing large extensions. Two such minimal proteins are *L. pneumophila* Lpp1663, which structure was solved by NMR (Immer et al. 2020), and *Neisseria meningitidis* minimal ProQ (NMB 1681), which structure was solved by X-ray crystallography (Chaulk et al. 2010). While the natural RNA ligands of Lpp1663 are not yet known, those of *N. meningitidis* minimal ProQ have been recently identified using CLIP-seq method (Bauriedl et al. 2020).

The 141-aa long *N. meningitidis* minimal ProQ protein (NMB 1681) consists mainly of the core FinO domain, and additionally has only short 19-aa long N-terminal, and 13-aa long C-terminal extensions (Chaulk et al. 2010; Olejniczak and Storz 2017). The comparison of the structures of the six copies of the protein present in crystallographic asymmetric unit showed that the 19-aa long N-terminal extension is likely flexible (Chaulk et al. 2010). Interestingly, while in the FinO domains of F-like plasmid FinO and *E. coli* ProQ there is a larger positively charged surface on the concave than on the convex face of the domain, in *N. meningitidis* minimal ProQ it extends on both the concave and the convex face (Chaulk et al. 2010; Olejniczak and Storz 2017).

Recent CLIP-seq study identified almost 200 mRNAs and sRNAs associated with *N. meningitidis* ProQ, in which the ProQ binding sites often overlapped intrinsic transcription terminators (Bauriedl et al. 2020). The direct binding of several of these RNAs to *N. meningitidis* ProQ was further supported by binding assays using purified components (Bauriedl et al. 2020). It was also previously shown that *N. meningitidis* ProQ bound tightly to the transcription terminator derived from FinP RNA (Chaulk et al. 2010).

To elucidate how the *N. meningitidis* minimal ProQ recognizes RNAs we compared the strength of ProQ binding to several of its natural RNA ligands using gelshift assay. Next we used mutant RNAs to determine what RNA features are essential for tight binding to *N. meningitidis* minimal ProQ. The data showed that the bottom part of the Rho-independent transcription terminator hairpin together with adjacent single-stranded sequences are recognized by the minimal ProQ, and that the 3ʹ terminal ribose has an important role in the interaction. The results of our studies suggest that although the minimal ProQ from *N. meningitidis* recognizes the same general RNA features as other FinO domain proteins, there are also subtle differences in RNA properties, which are optimal for binding by *N. meningitidis* ProQ in comparison with the isolated FinO domains from other proteins.

## RESULTS

To compare how tightly ProQ binds its different RNA ligands we selected a set of seven RNAs of different origins, which were previously identified in *N. meningitidis* using CLIP-seq method (Bauriedl et al. 2020) (Fig. 1). Among them were two sRNAs, Bns1 and AniS, two mRNA 5ʹ-UTRs, *carA*-5ʹ and *pnp*-5ʹ, two mRNA 3ʹ-UTRs, *rpmG*-3ʹ and *iga*-3ʹ, and an RNA originating from the intergenic region between the NMV_RS10770 and *app* genes (Fig. 1). Interestingly, AniS, which was the least enriched among ProQ ligands identified by CLIP-seq (Bauriedl et al. 2020), is also a ligand of *N. meningitidis* Hfq (Fantappie et al. 2011; Heidrich et al. 2017). The binding of Bns1 sRNA, AniS sRNA, the 5ʹ-UTR of *pnp*, and the 3ʹ-UTR of *rpmG* to purified *N. meningitidis* ProQ have already been shown (Bauriedl et al. 2020). The 3ʹ ends of sRNAs and mRNA 3ʹ-UTRs used in our binding studies were defined so as they contained the complete terminator structure, which overlapped with the reported CLIPseq peak (Bauriedl et al. 2020). The 3ʹ ends of mRNA 5-UTRs, *carA*-5ʹ and *pnp*-5ʹ, were defined by the 3ʹ end of the sequence of the reported CLIPseq peak (Bauriedl et al. 2020). The 5ʹ ends were defined to start with guanosine residues present in the natural sequence, or the guanosine residues were added to the 5ʹ ends, to ensure efficient *in vitro* transcription. Instead of full-length AniS sRNA, we used its 3ʹ-terminal fragment including the whole reported CLIPseq peak (Bauriedl et al. 2020), which we named AniS-3ʹ, and which bound ProQ with similar affinity as full-length AniS (Suppl. Fig. S1). In summary, the RNA molecules that we used included 90-nt long Bns1 sRNA, 55-nt long AniS-3ʹ, 65-nt long *pnp*-5ʹ, 64-nt long *carA-* 5ʹ, 48-nt long *rpmG*-3ʹ, 64-nt long *iga*-3ʹ, and 55-nt fragment of the intergenic region between the NMV_RS10770 and *app* genes, which we named *intergenic* (Fig. 1).

**Figure 1.**
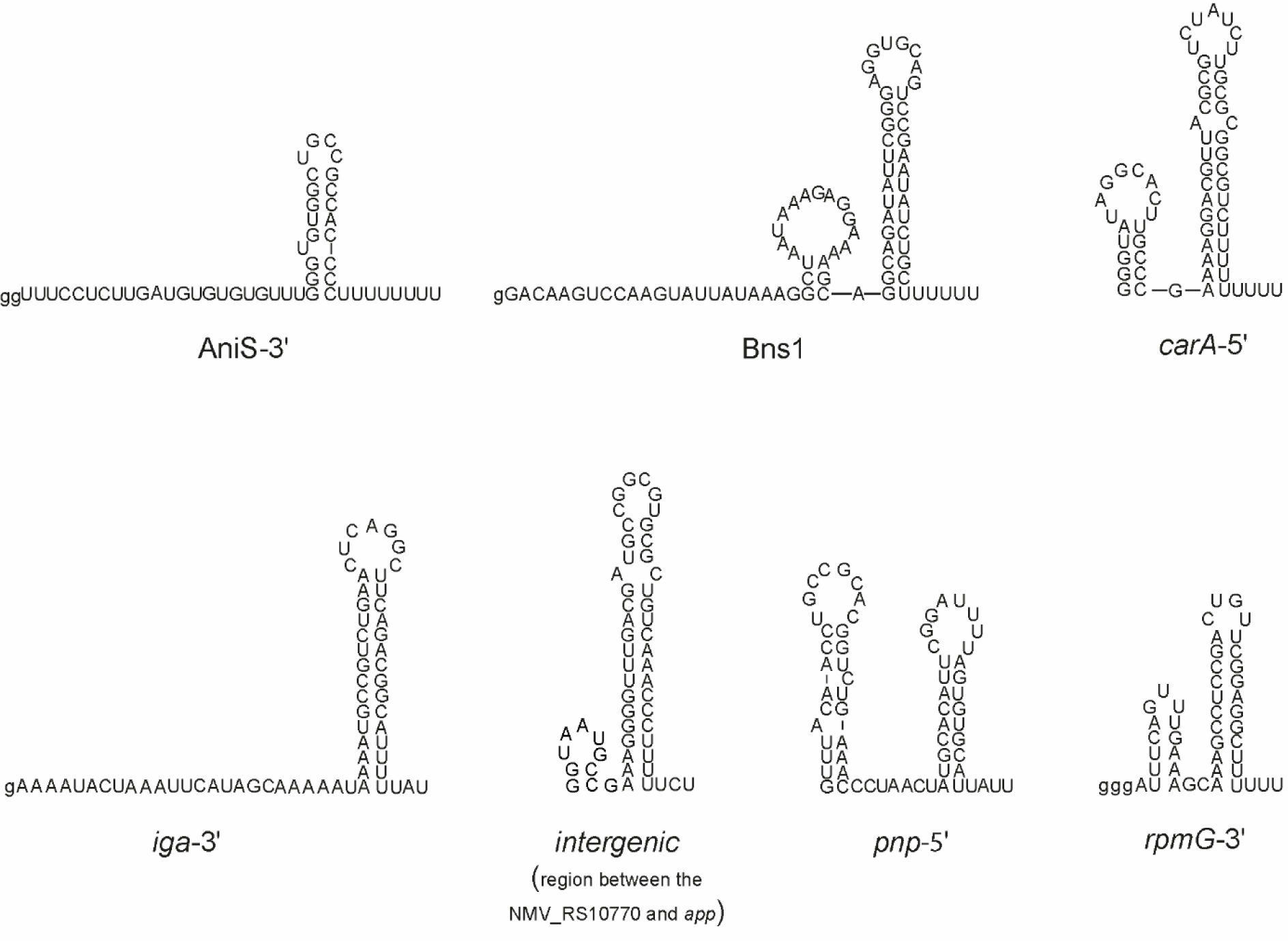
RNA molecules bound by ProQ protein in *Neisseria meningitidis*, which were used in this study. The secondary structures of RNAs AniS-3’, Bns1, *carA*-5’, *iga*-3’, *intergenic* (intergenic region between NMV_RS10770::*app*), *pnp*-5’ and *rpmG*-3’. The RNAs are selected from the list of the native *N. meningitidis* ProQ protein RNA ligands detected by CLIP-seq (Bauriedl et al. 2020). The lower case g denotes guanosine residue added on 5’ end to enable T7 RNA polymerase transcription. The RNA secondary structure predictions were performed in the *ViennaRNA* program (Lorenz et al. 2011).

We compared the binding affinities of the seven RNAs to ProQ using a gelshift assay (Fig. 2, Table 1, Suppl. Fig. S2). The data showed that all RNAs formed single complexes with ProQ in the studied concentration range (Fig. 2A, Suppl. Fig. S2). The binding of all RNAs to ProQ was tight with *K*_d_ values in low nanomolar range. The binding affinities of these RNAs for ProQ ranged from *K*_d_ value of 0.3 nM for *iga*-3ʹ to 7.8 nM for *pnp*-5ʹ. However, while the fraction bound of AniS-3ʹ, Bns1, *carA*-5ʹ, *intergenic*, and *rpmG*-3ʹ reached about 90% at saturation, that of *iga*-3ʹ and *pnp*-5ʹ saturated at only about 50 to 60% (Fig. 2, Table 1, Suppl. Fig. S2). This could suggest that these two RNAs formed alternative RNA conformations or that the complexes of these RNAs with ProQ partly dissociated during electrophoresis.

**Figure 2.**
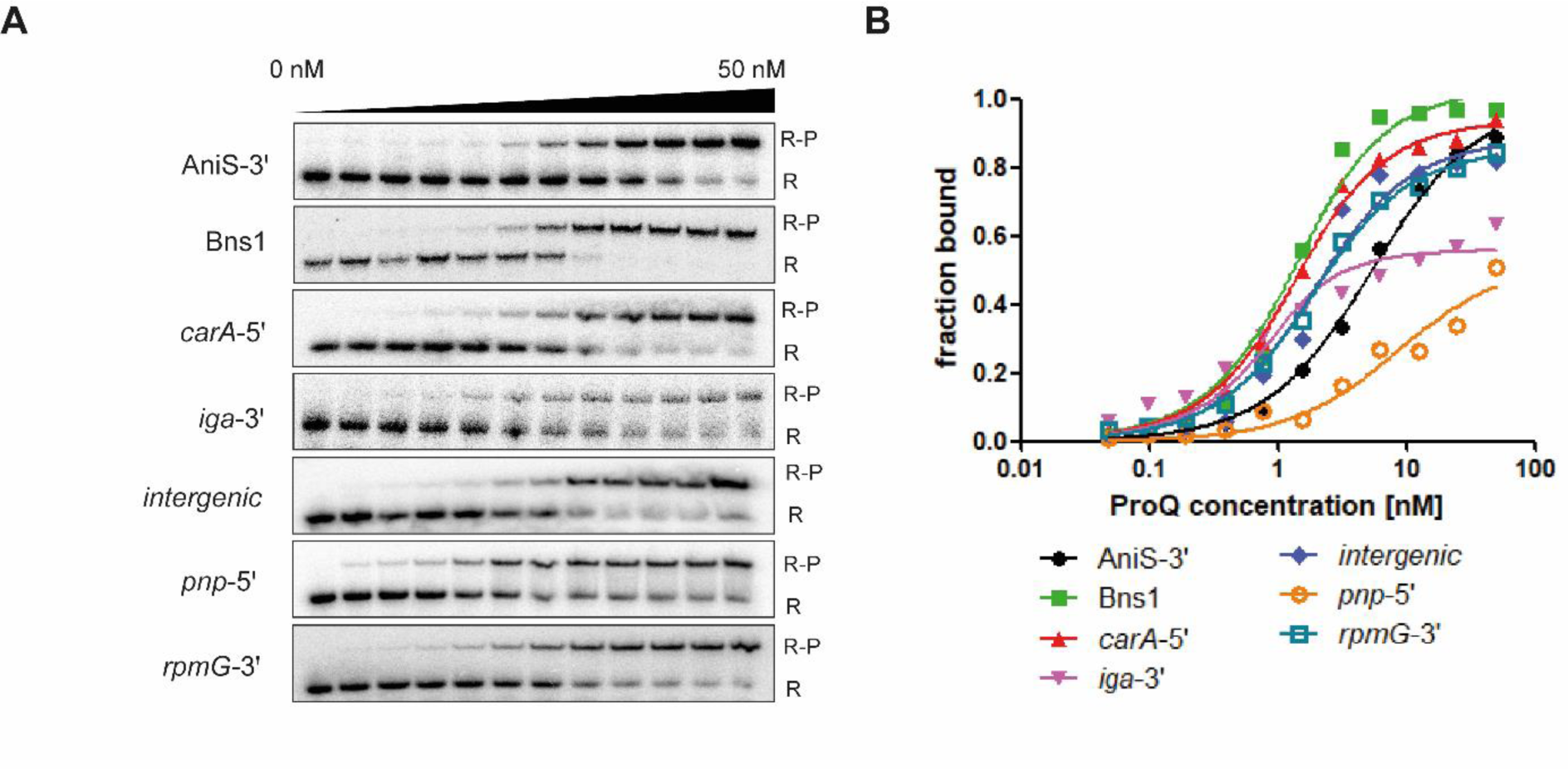
Equilibrium RNA binding to *Neisseria meningitidis* ProQ protein. (A) The binding of ^32^P-labeled RNAs AniS-3’, Bns1, *carA*-5’, *iga*-3’, *intergenic*, *pnp*-5’ and *rpmG*-3’ to ProQ was monitored using a gelshift assay. Free ^32^P-RNA is marked as R, RNA-ProQ complexes as R-P. (B) The fitting of the ProQ binding data from A using the quadratic equation provided *K*_d_ values of 4.9 nM for AniS-3’, 0.9 nM for Bns1, 0.9 nM for *carA*-5’, 0.3 nM for *iga*-3’, 1.4 nM for *intergenic,* 7.9 nM for *pnp*-5’ and 1.2 nM for *rpmG*-3’. The RNAs secondary structures predictions are presented in Fig. 1. Raw gel data for all RNAs are presented in Supplementary Figure S2. The average equilibrium dissociation constant (*K*_d_) values and the maximum RNA fraction bound calculated from at least three independent experiments are shown in Table 1.

**Table 1.** Equilibrium RNA binding to *N. meningitidis*.

| <sup>32</sup>P-RNA | *K<sub>d</sub>* [nM] (maximal fraction bound) |
| --- | --- |
| AniS-3' | 5 ± 0.4 (91 %) |
| Bns1 | 0.7 ± 0.5 (97 %) |
| *carA*-5' | 1.2 ± 0.4 (88 %) |
| *iga*-3' | 0.3 ± 0.3 (51 %) |
| *intergenic* | 1.4 ± 1.0 (86 %) |
| *pnp*-5' | 7.8 ± 0.8 (63 %) |
| *rpmG*-3' | 1 ± 0.5 (89 %) |

The *K<sub>d</sub>* values were obtained by fitting the data using the quadratic equation. The average *K<sub>d</sub>* values with standard deviations were calculated from at least three independent experiments.

Because it was previously observed that longer fragments of *pnp*-5ʹ and *rpmG*-3ʹ are also bound by ProQ (Bauriedl et al. 2020), we compared the binding of 3ʹ-extended versions of these two RNAs, which we named *pnp*-5ʹ-ext and *rpmG*-3ʹ-ext (Suppl. Figs. S3, S4). Both *pnp*-5ʹ-ext and *rpmG*-3ʹ-ext RNAs bound ProQ weaker than *pnp*-5ʹ and *rpmG*-3ʹ, respectively, because the fractions bound of each RNA at the maximum 50 nM concentration used were lower than 20%. Hence, extending the 3ʹ end of *rpmG*-3ʹ beyond the polyU tail of the terminator, or extending the 3ʹ end of *pnp*-5ʹ beyond a terminator-like structure, was detrimental for ProQ binding. This observation is consistent with previous reports that extending RNAs beyond their 3ʹ polyU tails was detrimental for RNA binding by the FinO domain of *E. coli* ProQ (Stein et al. 2020), and that RocR RNA with elongated 3ʹ tail bound less well to the FinO domain of *L. pneumophila* RocC protein (Kim et al. 2022).

Next, we analyzed how the length of the 3ʹ terminal polyU tail of the terminator affects the binding of *rpmG*-3ʹ and AniS-3ʹ RNAs to ProQ (Fig. 3, Suppl. Fig. S6). We focused on *rpmG*-3ʹ and AniS-3ʹ sRNA, because they markedly differed in the binding affinity to ProQ (Fig. 2, Table 1, Suppl. Fig. S2), and represented two important groups of RNA ligands of ProQ, mRNA 3ʹ-UTRs and sRNAs. For *rmpG*-3ʹ we compared eight variants, which differed with the length of the 3ʹ tail, including three which were longer than *rmpG*-3ʹ and included the following sequence of *rpmG* gene, while for AniS-3ʹ we compared seven length variants, of which AniS-3ʹ had the longest 8-uridine tail encoded in *aniS* gene.

**Figure 3.**
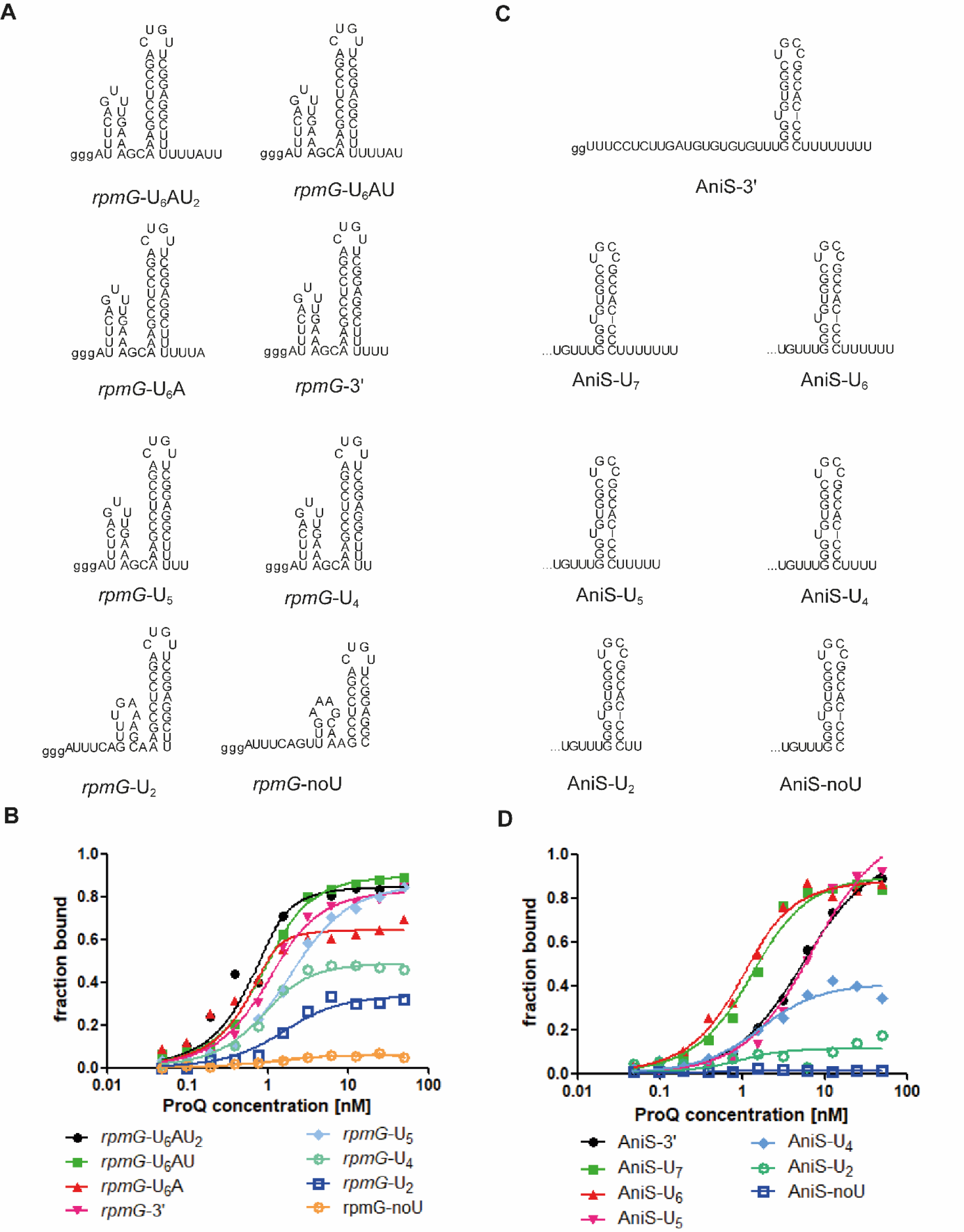
The 3’-terminal polyU tail is involved in *rpmG*-3’ and AniS-3’ binding to the *Neisseria meningitidis* ProQ protein. (A) *rpmG*-3’ constructs with different lengths of 3’ polyU tails. (B) The fitting of the ProQ binding data using the quadratic equation provided *K*_d_ values of 0.1 nM for *rpmG*-U_6_AU_2_, 0.4 nM for *rpmG*-U_6_AU, 0.1 nM for *rpmG*-U_6_A, 1.2 nM for *rpmG*-3’, 0.5 nM for *rpmG*-U_5_ and 0.4 nM for *rpmG*-U_4_, while the binding for *rpmG*-U_2_ reached saturation below 40 % of bound RNA fraction, and *rpmG*-noU did not reach saturation up to 50 nM concentration of the ProQ. (C) AniS-3’ constructs with different lengths of 3’ polyU tails. (D) The fitting of the ProQ binding data using the quadratic equation provided *K*_d_ values of 4.9 nM for AniS-3’, 0.8 nM for AniS-U_7_, 0.5 nM for AniS-U_6_, 6.6 nM for AniS-U_5_, 1.1 nM for AniS-U_4_, while the binding for AniS-U_2_ did not reach saturation up to 50 nM concentration of the ProQ. The binding of AniS-noU was essentially undetectable up to 50 nM concentration of the ProQ. The data in the plots for *rpmG*-3′ and AniS-3′ binding to ProQ are the same as in Figure 2. The lower case g denotes guanosine residue added on 5’ end to enable T7 RNA polymerase transcription. Gels corresponding to the data in the plots are shown in Supplementary Figure S6. The RNA secondary structure predictions were performed in the *ViennaRNA* program (Lorenz et al. 2011). The average equilibrium dissociation constant (*K*_d_) values and maximum RNA fraction bound calculated from at least three independent experiments are shown in Table 2.

**Table 2.** The length of the 3’ single-stranded tail following the Rho-independent terminator affects the binding of *rpmG*-3’ and AniS-3’ RNAs to *N. meningitidis* ProQ.

| <sup>32</sup>P-RNA | $K_d$ [nM] (maximal fraction bound) |
| --- | --- |
| *rpmG*-U<sub>6</sub>AU<sub>2</sub> | 0.2 ± 0.3 (88 %) |
| *rpmG*-U<sub>6</sub>AU | 0.2 ± 0.2 (89 %) |
| *rpmG*-U<sub>6</sub>A | < 0.5 (66 %) |
| *rpmG*-3' | 1 ± 0.5 (89 %)<sup>a</sup> |
| *rpmG*-U<sub>5</sub> | 0.5 ± 0.3 (85 %) |
| *rpmG*-U<sub>4</sub> | 2.6 ± 3.5 (58 %) |
| *rpmG*-U<sub>2</sub> | > 50 (27 %) |
| *rpmG*-noU | > 50 (6 %) |
| AniS-3' | 5 ± 0.4 (91 %)<sup>a</sup> |
| AniS-U<sub>7</sub> | 0.8 ± 0.2 (88 %) |
| AniS-U<sub>6</sub> | 0.7 ± 0.6 (79 %) |
| AniS-U<sub>5</sub> | 6.5 ± 1.3 (90 %) |
| AniS-U<sub>4</sub> | 0.9 ± 0.6 (53 %) |
| AniS-U<sub>2</sub> | > 50 (14 %) |
| AniS-noU | n. d. |

The $K_d$ values were obtained by fitting the data using the quadratic equation. The average $K_d$ values with standard deviations were calculated from at least three independent experiments.
<sup>a</sup> Data from Table 1.
n. d. - the binding was essentially undetected up to 50 nM concentration of the ProQ.

The data showed that the binding of *rpmG*-3ʹ and AniS-3ʹ to ProQ differently depended on the 3ʹ tail length (Fig. 3A,B, Table 2, Suppl. Fig. S6A). For *rmpG*-3ʹ the 3ʹ-tail lengths from 5 to 9 nucleotides ensured tight binding to ProQ. The binding of RNAs with 3ʹ tails of 4 or 2 nucleotides of length was markedly weaker, while the binding of that with no 3ʹ tail was barely detectable (Fig. 3A,B, Table 2, Suppl. Fig. S6A). Hence, even the *rmpG*-3ʹ mutant with the longest 9-nt tail bound tightly to ProQ. On the other hand, when the binding of RNAs derived from AniS-3ʹ was compared, the data showed that the derivatives with 3ʹ tails of 6 or 7 uridines of length, AniS-U_6_ and AniS-U_7_, bound tightest to ProQ (Fig. 3C,D, Table 2, Suppl. Fig. S6B). The RNA with the longest, 8-uridine 3ʹ tail, AniS-3ʹ, bound ProQ 5-fold weaker than AniS-U_6_. Additionally, shortening the 3ʹ tail below 6 uridines weakened the binding, with the exception of AniS-U_4_, which, however, had decreased maximum fraction bound. The binding of the shortest AniS-3ʹ derivatives, which had the 3ʹ tail of 2 uridines or no tail, was very weak or not detected (Fig. 3C,D, Table 2, Suppl. Fig. S6B). The observations that shortening of the 3ʹ tail of transcription terminator of *rmpG*-3ʹ below 5 uridines and that of AniS-3ʹ below 4 uridines weakens RNA binding to ProQ are similar to previous observations that shortening of the 3ʹ tail length below 4 uridines markedly weakened RNA binding to the FinO domain of *E. coli* ProQ (Stein et al. 2020), and that the 3ʹ tail of 3 uridines caused RocR RNA to bind much weaker to RocC protein than the 3ʹ tail of 5 uridines (Kim et al. 2022). Interestingly, the 3ʹ tail length of 9 nucleotides permits tight ProQ binding by *rmpG*-3ʹ, while the tail length of 8 nucleotides weakens the binding of AniS-3ʹ (Fig. 3, Table 2). Because the three uridines closest to the terminator of *rmpG*-3ʹ are involved in base-pairing it effectively shortens the single-stranded length of the 3ʹ tail. Hence, the differences between *rmpG*-3ʹ and AniS-3ʹ RNAs regarding the length of 3ʹ tail that is optimal for tight binding to *N. meningitidis* ProQ could be the result of the different involvement of their 3ʹ tails in RNA structure.

In the next step, we analyzed how the length of RNA sequence on the 5ʹ side of the terminator affects the binding of *rpmG*-3ʹ and AniS-3ʹ to ProQ (Fig. 4, Table 3, Suppl. Fig. S7). In these experiments we used chemically synthesized oligoribonucleotides. The longest *rpmG*-3ʹ-derived construct in these experiments was *rpmG*-45 which differed from *rpmG*-3ʹ by the absence of guanosine residues added to *rpmG*-3ʹ to enable efficient transcription. *rpmG*-45 bound ProQ somewhat weaker than *rpmG*-3ʹ but the binding achieved saturation at similar maximum fraction bound. Then a 10 nucleotide shorter construct was created, named *rpmG*-35. On the 5ʹ side of the G-C ending terminator hairpin this RNA had a 9-nt long sequence, which consisted of a 6-nt long single-stranded stretch and a 3-nt long stretch of adenosines base-paired with uridines of the 3ʹ tail. The affinity of *rpmG*-35 construct to ProQ had a similar *K*_d_ value as that of *rpmG*-45, which suggests that the 9-nt length of sequences consisting of single-stranded and douible-stranded stretches on the 5ʹ side of the terminator is sufficient for tight ProQ binding. However, when the 5ʹ part of the molecule was truncated further 2 nucleotides, the resulting *rpmG*-33 bound ProQ 3-fold weaker than *rpmG*-45. Further shortening resulted in *rpmG*-31, which had only two single-stranded residues on the 5ʹ end, and which bound ProQ very weakly. The binding of even more truncated *rpmG*-29 construct, which had the single-stranded portion of 5ʹ terminal RNA sequence completely removed, was also severely weakened as the binding to ProQ was not detected in the studied concentration range. Additionally, removing the 5ʹ terminal stretch of adenosine residues, which was base paired with uridines of 3ʹ tail, resulted in a construct, named *rpmG*-26, which binding to ProQ was also not detected.

**Figure 4.**
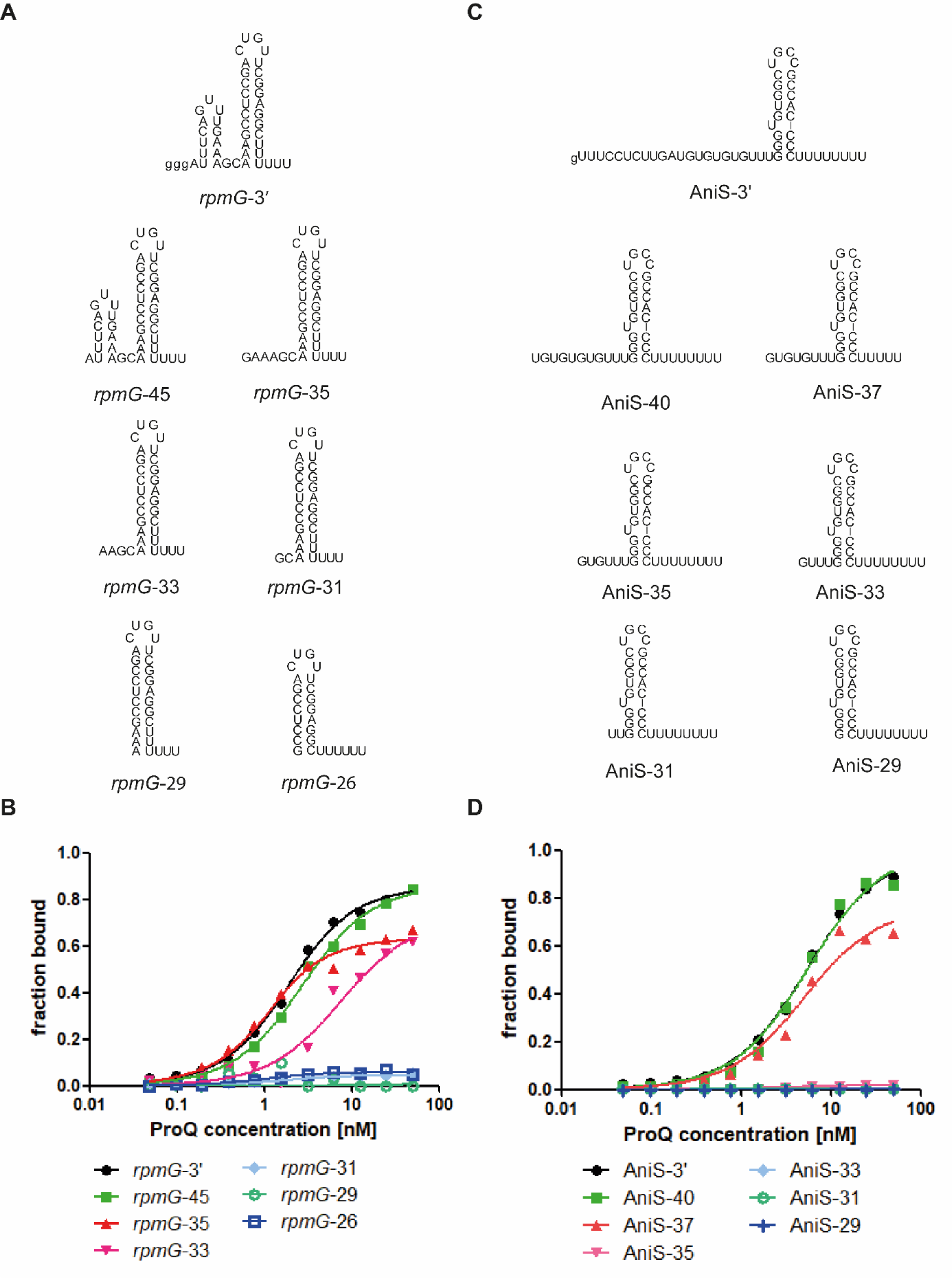
The 5’-terminal sequence preceding the terminator hairpin is involved *rpmG*-3’ and AniS-3’ binding to the *Neisseria meningitidis* ProQ protein. (A) *rpmG*-3’ constructs with different lengths of the 5’-terminal sequence. (B) The fitting of the ProQ binding data using the quadratic equation provided *K*_d_ values of 1.2 nM for *rpmG*-3’, 2.2 nM for *rpmG*-45, 0.6 nM for *rpmG*-35 and 7.0 nM for *rpmG*-33, while the binding for *rpmG*-31 did not reach saturation up to 50 nM concentration of the ProQ. The binding of *rpmG*-29 and *rpmG*-26 was essentially undetectable up to 50 nM concentration of the ProQ. (C) AniS-3’ constructs with different lengths of the 5’-terminal sequence. (D) The fitting of the ProQ binding data using the quadratic equation provided *K*_d_ values of 4.9 nM for AniS-3’, 5.0 nM for AniS-40, 4.5 nM for AniS-37, while the binding of AniS-35 was barely detected, and that of AniS-33, AniS-31 and AniS-29 was essentially undetected up to 50 nM concentration of the ProQ. The data in the plots for *rpmG*-3′ and AniS-3′ binding to ProQ are the same as in Figure 2. The lower case g denotes guanosine residue added on 5’ end to enable T7 RNA polymerase transcription. Gels corresponding to the data in the plots are shown in Supplementary Figure S7. The RNA secondary structure predictions were performed in the *ViennaRNA* program (Lorenz et al. 2011). The average equilibrium dissociation constant (*K*_d_) values and maximum RNA fraction bound calculated from at least three independent experiments are shown in Table 3.

**Table 3.** The length of the sequence on the 5’ side of the Rho-independent terminator affects the binding of *rpmG*-3’ and AniS-3’ RNAs to *N. meningitidis* ProQ.

| <sup>32</sup>P-RNA | $K_d$ [nM] (maximal fraction bound) |
| --- | --- |
| *rpmG*-3' | $1 \pm 0.5$ (89 %)<sup>a</sup> |
| *rpmG*-45 | $2.8 \pm 1.7$ (90 %) |
| *rpmG*-35 | $0.8 \pm 0.2$ (71 %) |
| *rpmG*-33 | $7.3 \pm 2.5$ (63 %) |
| *rpmG*-31 | > 50 (8 %) |
| *rpmG*-29 | n. d. |
| *rpmG*-26 | n. d. |
| AniS-3' | $5 \pm 0.4$ (91 %)<sup>a</sup> |
| AniS-40 | $3.9 \pm 4.4$ (83 %) |
| AniS-37 | $4.0 \pm 1.2$ (60 %) |
| AniS-35 | > 50 (2 %) |
| AniS-33 | n. d. |
| AniS-31 | n. d. |
| AniS-29 | n. d. |

The $K_d$ values were obtained by fitting the data using the quadratic equation. The average $K_d$ values with standard deviations were calculated from at least three independent experiments.
<sup>a</sup> Data from Table 1.
n. d. - the binding was essentially undetected up to 50 nM concentration of the ProQ.

When the 5ʹ truncated constructs of AniS-3ʹ were analyzed we also observed dependence of ProQ binding affinity on the length of the 5ʹ terminal single-stranded sequence. When the 5ʹ terminal single-stranded region was shortened to 11 nucleotides in the AniS-40 construct it did not markedly affect the binding affinity as the *K*_d_ value was similar as that of AniS-3ʹ. Of note, AniS-3ʹ does not contain an adenosine stretch bordering with the terminator. Hence, the whole sequence 5ʹ terminal to the G-C closing base pair of the terminator is single-stranded. Further truncation to 8 nucleotide residues resulted in AniS-37, which had the same *K*_d_ value of ProQ binding, but the maximum fraction bound was lower. However, when the 5ʹ terminal single-stranded sequence was shortened to 6 nucleotides in the AniS-35 construct the binding to ProQ was abolished, and the same effect was observed when this sequence was shortened to 4, and 2 nucleotides or the whole sequence was removed, in AniS-33, AniS-31, and AniS-29, respectively. Hence, for the efficient binding of AniS-3ʹ to ProQ the length of at least eight nucleotides of single-stranded sequence on the 5ʹ side of the terminator is necessary. The importance of the single-stranded sequence on the 5ʹ side of the terminator for RNA binding to ProQ is consistent with previous observation that a seven-nucleotide long 5ʹ terminal single-stranded sequence was sufficient for tight binding of a model RNA construct derived from *cspE*-3ʹ to *E. coli* ProQ, while the complete removal of 5ʹ terminal single-stranded sequence abolished the binding (Stein et al. 2023). In further support of the importance of the sequence 5ʹ of the terminator for *N. meningitidis* ProQ binding, it was previously shown that the CLIP-seq peak of AniS included also the sequence on the 5ʹ side of the terminator (Bauriedl et al. 2020). In summary, our data showed that a longer single-stranded sequence on the 5ʹ side was necessary for tight binding of AniS-3ʹ than *rpmG*-3ʹ.

To test what is the contribution of the double-stranded portion of the terminator hairpin to RNA binding of ProQ we designed variants of *rpmG*-3ʹ and AniS-3ʹ, with gradually shortened terminator hairpin stems (Fig. 5, Table 4, Suppl. Fig. S8). At first we replaced the apical loop of the terminator hairpin of *rpmG*-3ʹ with GAAA tetraloop to increase the stability of the shortened hairpin (*rpmG*-loop). Then, we gradually shortened the hairpin stem in two-base-pair steps, thus creating *rpmG*-loop-44, *rpmG*-loop-40 and *rpmG*-loop-36 (Fig. 5A). We did not design a shorter construct, because it was predicted by *ViennaRNA* software not to retain a hairpin structure. The data showed that all mutants, including *rpmG*-loop-36 with the shortest hairpin, which consisted of 2 G-C base pairs followed by 3 A-U base pairs, bound ProQ no weaker than *rpmG*-loop with natural length of hairpin stem (Fig. 5B, Table 4, Suppl. Fig. S8A). This result suggests that *N. meningitidis* ProQ binds the lower part of the terminator hairpin of *rpmG*-3ʹ.

**Figure 5.**
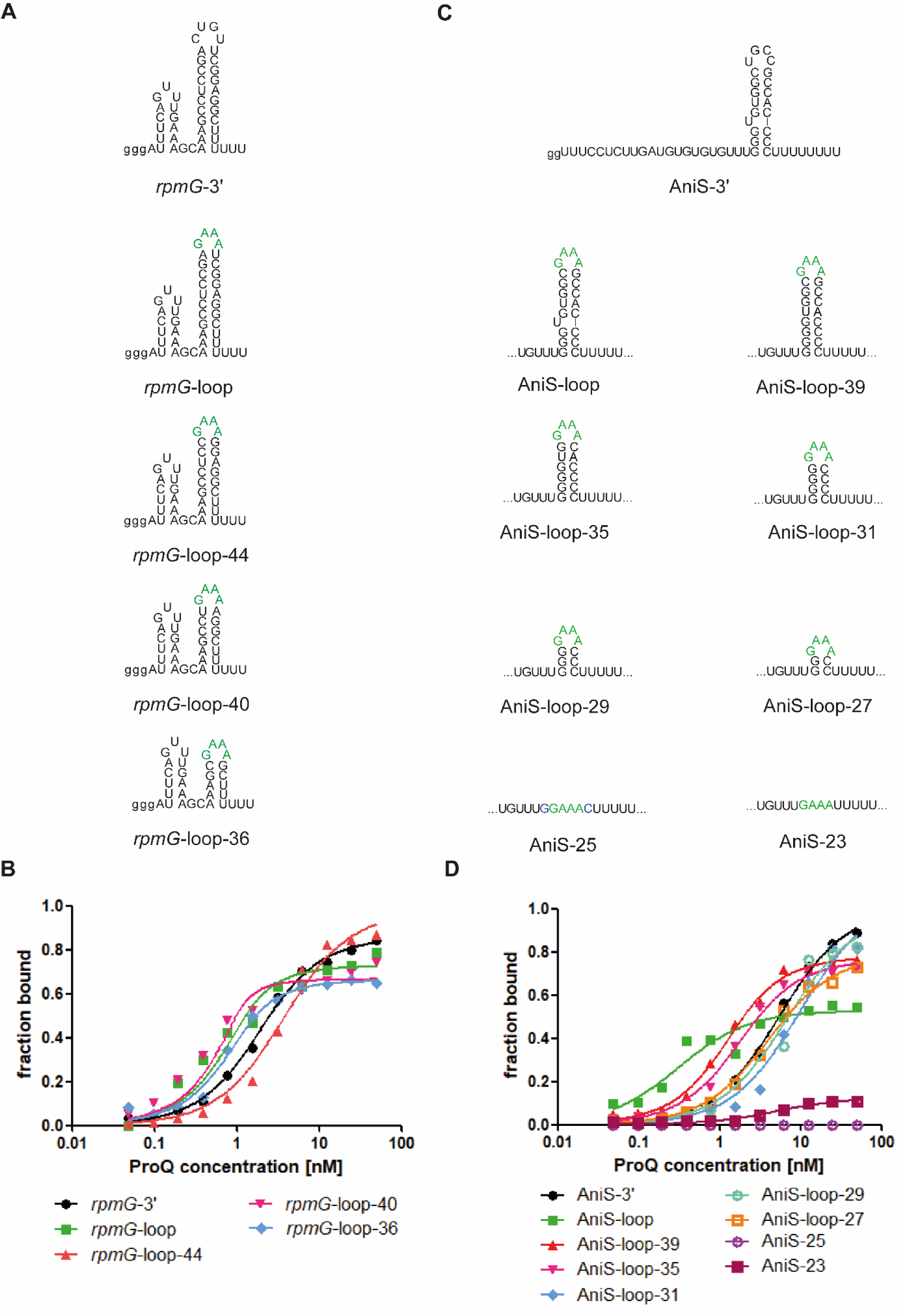
The lower part of the terminator hairpin is involved in *rpmG*-3’ and AniS-3’ binding to the *Neisseria meningitidis* ProQ protein. (A) *rpmG*-3’ mutants with shorter terminator stems were constructed by replacement of the native apical loop CUGU with the tetraloop GAAA and gradual removal of base-pairs from the top of the terminator stem. (B) The fitting of the ProQ binding data using the quadratic equation provided *K*_d_ values of 1.2 nM for *rpmG*-3’, 0.3 nM for *rpmG*-loop, 3.2 nM for *rpmG*-loop-44, 0.6 nM for *rpmG*-loop-40 and 0.3 nM for *rpmG*-loop-36. (C) AniS-3’ mutants with shorter terminator stems were constructed by replacement of the native apical loop UGCC with the tetraloop GAAA, removing the bulge, and gradual removal of base-pairs from the top of the terminator stem. (D) The fitting of the ProQ binding data using the quadratic equation provided *K*_d_ values of 4.9 nM for AniS-3’, 0.1 nM for AniS-loop, 0.7 nM for AniS-loop-39, 1.3 nM for AniS-loop-35, 9.2 nM for AniS-loop-31, 6.2 nM for AniS-loop-29, 3.5 nM for AniS-loop-27, while the binding for AniS-25 was barely detected, and AniS-23 was essentially undetected up to 50 nM concentration of the ProQ. The data in the plots for *rpmG*-3′ and AniS-3′ binding to ProQ are the same as in Figure 2. The lower case g denotes guanosine residue added on 5’ end to enable T7 RNA polymerase transcription. Green font indicates GAAA tetraloop. Gels corresponding to the data in the plots are shown in Supplementary Figure S8. The RNA secondary structure predictions were performed in the *ViennaRNA* program (Lorenz et al. 2011). The average equilibrium dissociation constant (*K*_d_) values and maximum RNA fraction bound calculated from at least three independent experiments are shown in Table 4.

**Table 4.** The bottom part of the terminator stem of *rpmG*-3’ and AniS-3’ RNAs is recognized by *N. meningitidis* ProQ.

| <sup>32</sup>P-RNA | $K_d$ [nM] (maximal fraction bound) |
| --- | --- |
| *rpmG*-3' | $1 \pm 0.5$ (89 %)<sup>a</sup> |
| *rpmG*-loop | $0.5 \pm 0.3$ (81 %) |
| *rpmG*-loop-44 | $4 \pm 2$ (91 %) |
| *rpmG*-loop-40 | $1.2 \pm 1.5$ (80 %) |
| *rpmG*-loop-36 | $0.1 \pm 0.2$ (64 %) |
| AniS-3' | $5 \pm 0.4$ (91 %)<sup>a</sup> |
| AniS-loop | $0.4 \pm 0.2$ (53 %) |
| AniS-loop-39 | $1.1 \pm 1.1$ (72 %) |
| AniS-loop-35 | $1 \pm 0.3$ (75 %) |
| AniS-loop-31 | $8.3 \pm 3.7$ (82 %) |
| AniS-loop-29 | $5.8 \pm 3.9$ (81 %) |
| AniS-loop-27 | $4.4 \pm 2.8$ (62 %) |
| AniS-25 | > 50 % (9 %) |
| AniS-23 | n. d. |

The $K_d$ values were obtained by fitting the data using the quadratic equation, except for the data for AniS-loop construct, which were fit using the equation for one site-specific binding with Hill slope model. The average $K_d$ values with standard deviations were calculated from at least three independent experiments.
<sup>a</sup> Data from Table 1.
n. d. - the binding was essentially undetected up to 50 nM concentration of the ProQ.

Next, we designed the constructs of AniS-3ʹ sRNA with shortened terminator stems (Fig. 5C). In this series of molecules the apical loop of AniS-3ʹ was also replaced with GAAA tetraloop. Additionally, we removed the single-uridine bulge located above the third base pair of the hairpin stem, thus creating a construct, which we named AniS-loop-39. Because this construct bound tightly to ProQ, and had a continuous double-stranded stem (Fig. 5D, Table 4, Suppl. Fig. S8B), we then designed truncated constructs based on AniS-loop-39. The data showed that shortening this construct by two base pairs to AniS-loop-35, which had a 6-base-pair stem did not weaken the binding. On the other hand, further shortening in 2-base-pair increments to AniS-loop-31, AniS-loop-29, and AniS-loop-27, which had the shortest stem consisting of only two G-C base pairs, resulted in at least 4-fold weaker binding in comparison to AniS-loop-39. However, even AniS-loop-27 bound tightly to ProQ with nanomolar *K*_d_. Complete removal of the hairpin stem in AniS-loop-25 and AniS-loop-23 completely abolished the binding. This suggests that the lowest two G-C base pairs are the part of the terminator hairpin of AniS-3ʹ which is essential for the binding to ProQ. The involvement of the lower parts of the terminator hairpins of *rpmG*-3ʹ and AniS-3ʹ in binding to ProQ is consistent with previous observations that this region is important for RNA binding by other FinO domain proteins (Arthur et al. 2011; Stein et al. 2020; Kim et al. 2022).

To better understand how the terminator hairpin is recognized by ProQ, we explored what is the contribution of the terminal residue of the RNA 3ʹ tail to its binding by ProQ (Fig. 6, Table 5, Suppl. Fig. S9). To achieve that we designed RNA constructs derived from *rpmG*-3ʹ and AniS-3ʹ, in which the 3ʹ terminal residue was modified in a way that could affect hydrogen bonding interactions. The modified RNAs were chemically synthesized. As the model RNAs we selected the constructs *rpmG*-35 and AniS-37 (Fig. 4A,C), because they are shorter than *rpmG*-3ʹ and AniS-3ʹ, respectively, but retain the ability to bind tightly to ProQ (Table 3). Both these molecules have uridines as the 3ʹ terminal residue. We designed two derivatives of each of these RNAs with modifications of the 3ʹ terminal uridine. One of them had a 2ʹ-deoxyribose, and the other one had the 3ʹ-OH group phosphorylated. Additionally, for both RNAs we designed a derivative, in which the 3ʹ terminal uridine was replaced with cytidine, and another one, in which it was replaced with 2ʹ,3ʹ-dideoxycytidine.

**Figure 6.**
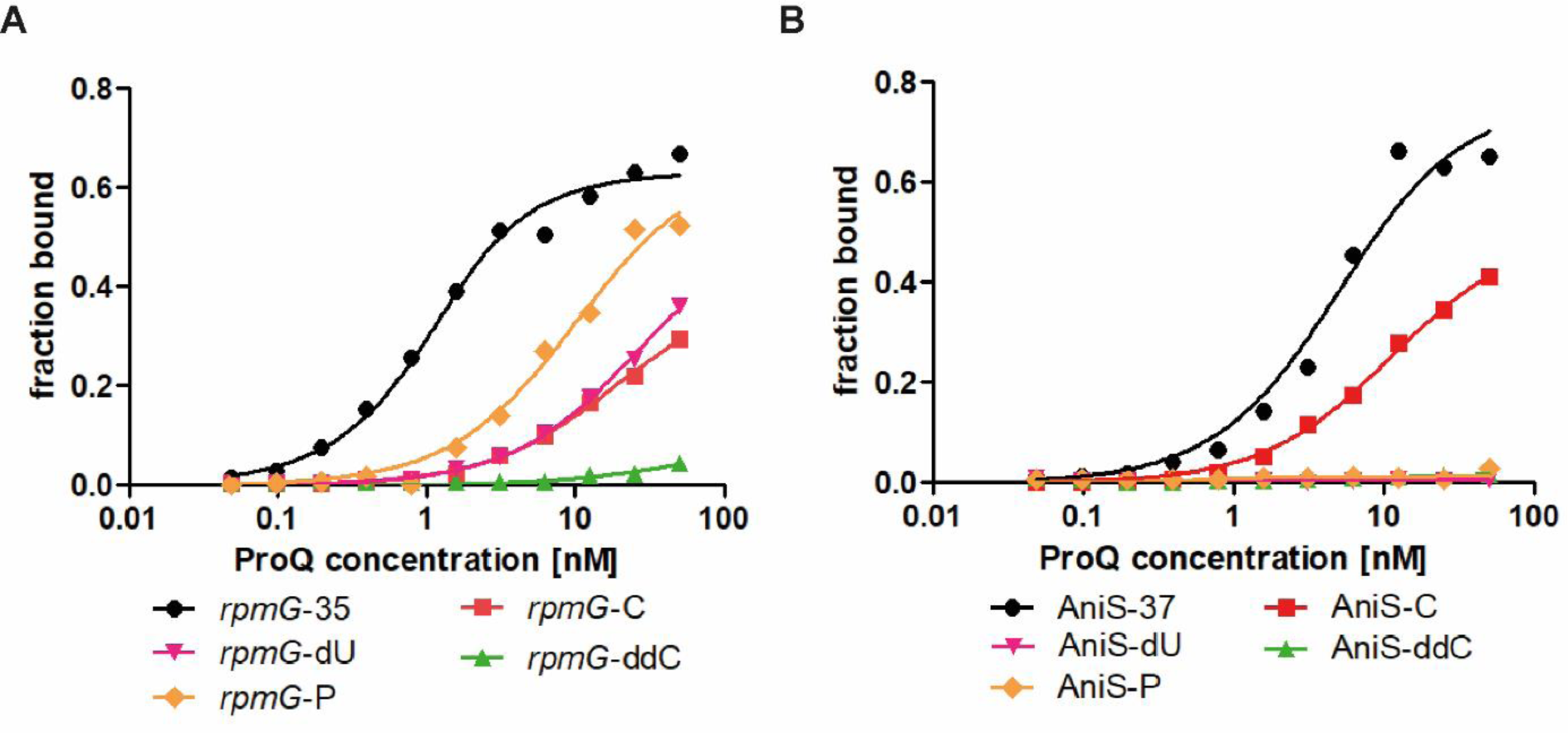
3ʹ-terminal uridine is specifically recognized through the ribose 2ʹ- and 3ʹ-OH groups when RNA is bound by the *Neisseria meningitidis* ProQ protein. (A) The fitting of the ProQ binding data using the quadratic equation provided *K*_d_ values of 0.6 nM for *rpmG*-35, while the binding of *rpmG*-dU, *rpmG*-P and *rpmG*-C did not reach saturation up to 50 nM concentration of the ProQ. The binding of *rpmG*-ddC was essentially undetectable up to 50 nM concentration of the ProQ. (B) The fitting of the ProQ binding data using the quadratic equation provided *K*_d_ values of 4.5 nM for AniS-37 and 10.5 nM for AniS-C, while the binding for AniS-dU, AniS-P and AniS-ddC was essentially undetectable up to 50 nM concentration of the ProQ. The data in the plots for *rpmG*-35 and AniS-37 binding to ProQ are the same as in Figure 4. Gels corresponding to the data in the plots are shown in Supplementary Figure S9. The average equilibrium dissociation constant (*K*_d_) values and maximum RNA fraction bound calculated from at least three independent experiments are shown in Table 5.

**Table 5.** The 2’ and 3’ hydroxyl groups of the 3’-terminal nucleoside are important for RNA recognition by *N. meningitidis* ProQ.

| <sup>32</sup>P-RNA | $K_d$ [nM] (maximal fraction bound) |
| --- | --- |
| *rpmG*-35 | $0.8 \pm 0.2$ (71 %)<sup>a</sup> |
| *rpmG*-dU | > 50 (35 %) |
| *rpmG*-P | > 50 (37 %) |
| *rpmG*-C | > 50 (23 %) |
| *rpmG*-ddC | n. d. |
| AniS-37 | $4.0 \pm 1.2$ (60 %)<sup>a</sup> |
| AniS-dU | n. d. |
| AniS-P | n. d. |
| AniS-C | $11 \pm 6.9$ (43 %) |
| AniS-ddC | n. d. |

The $K_d$ values were obtained by fitting the data using the quadratic equation. The average $K_d$ values with standard deviations were calculated from at least three independent experiments.
<sup>a</sup> Data from Table 3.
n. d. - the binding was essentially undetected up to 50 nM concentration of the ProQ.

When the binding of 3ʹ-terminally modified constructs of *rpmG*-35 was compared the data showed that all modifications weakened RNA binding to ProQ (Fig. 6A, Table 5, Suppl. Fig. S9A). When the 3ʹ-OH group of terminal uridine was blocked by phosphorylation it caused at least 10-fold weaker binding of *rpmG*-P in comparison to unmodified *rpmG*-35. However, removing the 2ʹ-OH group by replacing ribose with 2ʹ deoxyribose in *rpmG*-dU construct resulted in even stronger detrimental effect on binding than that of *rpmG*-P (Fig. 6A). A similar weakening of binding to ProQ was observed when the 3ʹ terminal uridine was replaced with cytidine in *rpmG*-C. On the other hand, removing both 2ʹ-OH and 3ʹ-OH group of the terminal cytidine in *rpmG*-ddC completely abolished its binding to ProQ. The effects of modifications on the binding of AniS-37 derived constructs were even stronger, because the binding of the three constructs, AniS-dU, AniS-P, and AniS-ddC, in which the 2ʹ-OH and/or 3ʹ-OH groups were modified, could not be detected (Fig. 6B, Table 5, Suppl. Fig. S9B). On the other hand, substituting the 3ʹ terminal uridine with cytidine had only a 2-fold detrimental effect on binding. The observation that 2ʹ-OH and 3ʹ-OH groups of the 3ʹ terminal ribose are important for RNA binding by ProQ is consistent with previous studies on the effects of modifying these groups on RNA binding by F-like plasmid FinO protein and *L. pneumophila* RocC (Arthur et al. 2011; Kim et al. 2022). The co-crystal structure of *L. pneumophila* RocC explains these effects by showing that that 2ʹ-OH and 3ʹ-OH groups of RocR RNA are within hydrogen bonding distance to conserved amino acids of RocC protein (Kim et al. 2022). On the other hand, no direct contacts were observed in the crystal structure between the uracil base of the 3ʹ terminal nucleoside and RocC protein (Kim et al. 2022), which is consistent with small effect of replacing uridine to cytidine in AniS-37 (Fig. 6, Table 5, Suppl. Fig. S9). Hence, we hypothesize that the negative effect of cytidine substitution in *rpmG*-C could be a result of changes in local base-pairing involving the cytidine rather than the disruption of specific contacts with *N. meningitidis* ProQ.

Because it was previously observed that RNAs bound by *E. coli* ProQ often had A-rich motifs on the 5ʹ side of the terminators (Stein et al. 2020), we used *WebLogo* software to compare the nucleotide content of the 10-nt long sequence on the 5ʹ side of the terminator in top 40 previously identified RNA ligands of *N. meningitidis* ProQ, in which the CLIP-seq peak overlapped with intrinsic transcription terminators (Bauriedl et al. 2020) (Suppl. Fig. S4, Suppl. Table S4). As controls, we also compared the nucleotide contents of the corresponding sequences in top 40 3ʹ-UTRs and sRNAs identified as ligands of *N. meningitidis* Hfq using RIP-seq (Heidrich et al. 2017), and in randomly selected 98 transcripts of *N. meningitidis* transcriptome. The analysis showed that there were no statistically significant differences between these three data sets, and in all of them a short sequence on the 5ʹ side of the terminator was enriched in adenosines (Suppl. Fig. S4, Suppl. Table S4). This shows that the A-enrichment on the 5ʹ side of the terminator is a general feature of *N. meningitidis* transcriptome. On the other hand, very few RNA ligands of Hfq, including AniS, had U-rich sequence motifs in this region (Suppl. Table S4).

To test if the nucleotide content of the sequence immediately 5ʹ of the terminator hairpin affects RNA binding to *N. meningitidis* ProQ we introduced substitutions in this region in *rpmG*-3ʹ and AniS-3ʹ (Fig. 7, Table 6, Suppl. Fig. S10). While *rpmG*-3ʹ has a three-adenosine stretch opposite to its 3ʹ tail, AniS-3ʹ has a three uridine stretch in the corresponding position (Fig. 1). To explore the importance of such a motif for ProQ binding we designed two kinds of constructs. In *rpmG*-3ʹ we replaced either 2 or 3 adenosines of the A-rich motif by uridines, in this way creating *rpmG*-2AtoU and *rpmG*-3AtoU. On the other hand, in AniS-3ʹ, we substituted either 2 or 3 uridines present on the 5ʹ side of AniS-3ʹ terminator by adenosines, thus creating AniS-2UtoA and AniS-3UtoA constructs. The data showed that substituting adenosines with uridines moderately weakened the ProQ binding of *rpmG*-2AtoU and *rpmG*-3AtoU, because the *K*_d_ values were about 2-fold or 4-fold weaker, respectively, for each construct (Fig. 7B, Table 6, Suppl. Fig. S10A). On the other hand, the substitutions of uridines to adenosines in corresponding positions of AniS-3ʹ strengthened the binding of AniS-2UtoA and AniS-3UtoA, because the *K*_d_ values were either 2-fold or more than 10-fold tighter, respectively, for each construct (Fig. 7D, Table 6, Suppl. Fig. S10B).

**Figure 7.**
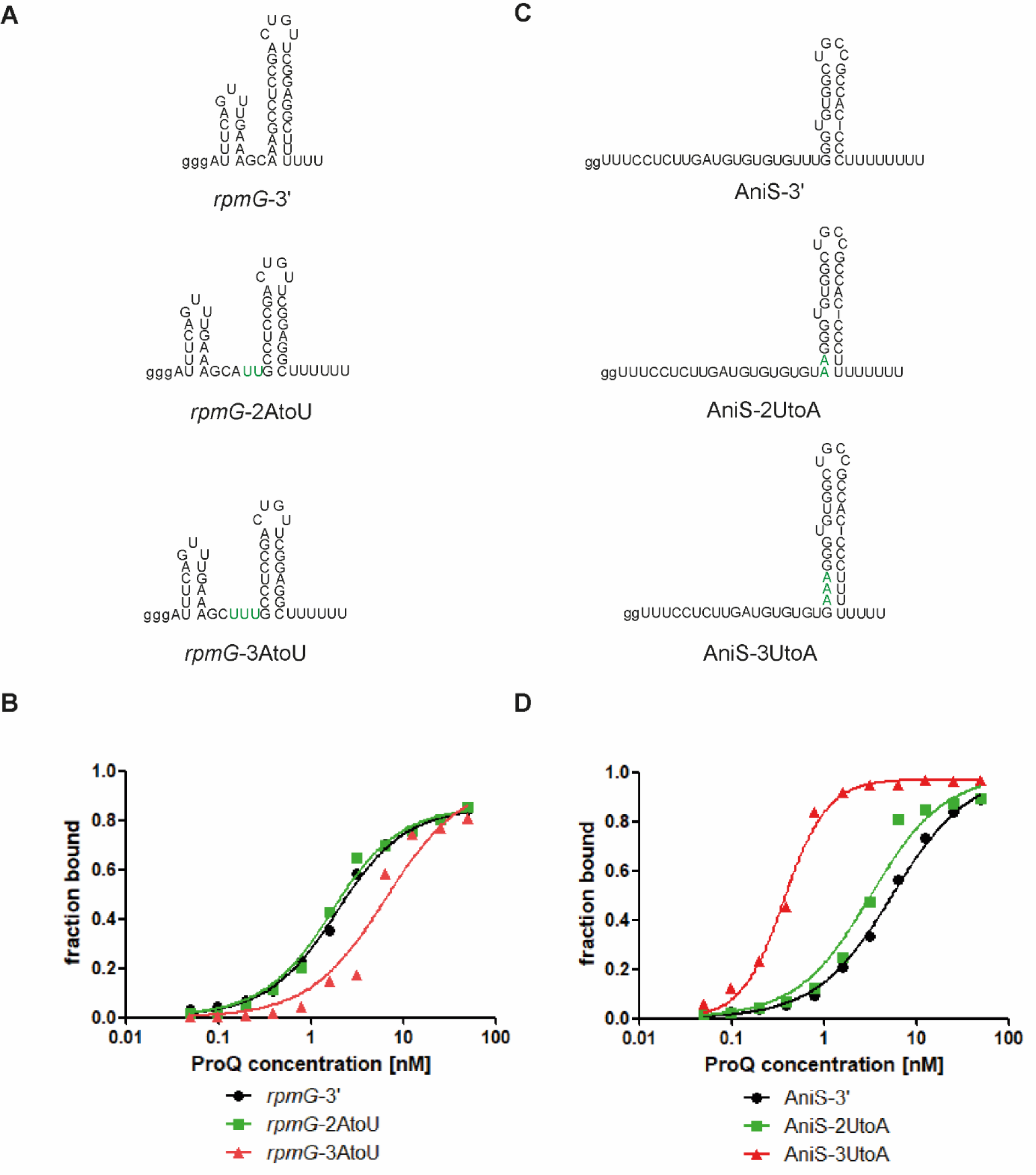
RNA mutations in the sequence at the 5’ side of the terminator stem affect the *rpmG*-3’ and AniS-3’ binding to *Neisseria meningitidis* ProQ protein. (A) Secondary structures of *rpmG*-3’ and its mutants. (B) The fitting of the ProQ binding data using the quadratic equation provided *K*_d_ values of 1.2 nM for *rpmG*-3’, 1.1 nM for *rpmG*-2AtoU and 5.8 nM for *rpmG*-3AtoU. (C) Secondary structures of AniS-3’ and its mutants. (D) The fitting of the ProQ binding data using the quadratic equation provided *K*_d_ values of 4.9 nM for AniS-3’ and 2.6 nM for AniS-2UtoA. The fitting of AniS-3UtoA to the equation for one site-specific binding with Hill slope model provided *K*_d_ value of 0.3 nM. The data in the plots for *rpmG*-3′ and AniS-3′ binding to ProQ are the same as in Figure 2. The lower case g denotes guanosine residue added on 5’ end to enable T7 RNA polymerase transcription. Green font indicates the introduced substitutions. Gels corresponding to the data in the plots are shown in Supplementary Figure S10. The RNA secondary structure predictions were performed in the *ViennaRNA* program (Lorenz et al. 2011). The average equilibrium dissociation constant (*K*_d_) values and maximum RNA fraction bound calculated from at least three independent experiments are shown in Table 6.

**Table 6.** Mutations in the sequence on the 5’ side of the terminator stem affect the binding of the *rpmG*-3’ and AniS-3’ to *N. meningitidis* ProQ.

| <sup>32</sup>P-RNA | *K<sub>d</sub>* [nM] (maximal fraction bound) |
| --- | --- |
| *rpmG*-3' | 1 ± 0.5 (89 %) <sup>a</sup> |
| *rpmG*-2AtoU | 1.7 ± 0.8 (91 %) |
| *rpmG*-3AtoU | 4.2 ± 2.7 (78 %) |
| AniS-3' | 5 ± 0.4 (91 %) <sup>a</sup> |
| AniS-2UtoA | 2.8 ± 0.6 (88 %) |
| AniS-3UtoA | 0.3 ± 0.1 (97 %) |

The *K<sub>d</sub>* values were obtained by fitting the data using the quadratic equation, except the data for AniS-3UtoA which was fit using the equation for one site-specific binding with Hill slope model. The average *K<sub>d</sub>* values with standard deviations were calculated from at least three independent experiments.
<sup>a</sup> Data from Table 1.

## DISCUSSION

The data presented here showed that the minimal ProQ protein from *N. meningitidis* recognizes three distinct parts of the intrinsic transcription terminator structures, which are the 5ʹ adjacent single-stranded sequence, the lower part of the terminator hairpin and the 3ʹ terminal tail (Figs. 3-5, Tables 2-4, Suppl. Fig. S6-S8). The same regions of the terminator are also important for RNA binding by other FinO-domain proteins including the F-like plasmid FinO protein, and the FinO domains of *E. coli* ProQ and *L. pneumophila* RocC proteins (Jerome and Frost 1999; Arthur et al. 2011; Stein et al. 2020; Kim et al. 2022; Stein et al. 2023).

The single-stranded sequence on the 5ʹ side of the terminator is essential for tight binding of *rpmG*-3ʹ and AniS-3ʹ to *N. meningitidis* ProQ (Fig. 4, Table 3, Suppl. Fig. S7), while it has varied contributions to RNA binding by other FinO domain proteins (Jerome and Frost 1999; Attaiech et al. 2016; Kim et al. 2022; Stein et al. 2023). The truncation of the 5ʹ sequence to 5 nucleotides or less in *rpmG*-3ʹ (including 2 single-stranded and 3 double-stranded nucleotides) or to 6 single-stranded nucleotides or less in AniS-3ʹ abolished their binding to *N. meningitidis* ProQ (Fig. 4, Table 3, Suppl. Fig. S7). The importance of the single-stranded sequence on the 5ʹ side of the transcription terminator was also reported for the binding of other FinO domain proteins. A 4-nt long single-stranded sequence on the 5ʹ side of terminator in a fragment of FinP RNA contributed to the strength of this RNA binding to F-like plasmid FinO protein, and when transferred on another hairpin it also improved its binding by FinO (Jerome and Frost 1999). However, in contrast to observations for AniS-3ʹ and *rpmG*-3ʹ binding to *N. meningitidis* ProQ (Fig. 4, Table 3, Suppl. Fig. S7), the complete removal of the 5ʹ-adjacent sequence from the terminator of FinP only moderately affected the FinO binding (Jerome and Frost 1999). Additionally, a short 1-nt sequence on the 5ʹ side of the terminator was sufficient for the tight binding of a fragment of RocR RNA to the FinO domain of *L. pneumophila* RocC protein (Attaiech et al. 2016; Kim et al. 2022). On the other hand, the binding of a model RNA derived from *cspE*-3ʹ to *E. coli* ProQ was abolished when the 5ʹ sequence was truncated to only 4 double-stranded nucleotides remaining or when it was completely removed (Stein et al. 2023). These data suggest that RNA binding by both *N. meningitidis* ProQ and *E.coli* ProQ is strongly dependent on the single-stranded sequence on the 5ʹ side of the terminator, while it has smaller contributions to RNA binding by the F-like plasmid FinO and *L. pneumophila* RocC proteins.

The data presented here and the previous studies of several FinO-domain proteins indicate that the lower part of the terminator hairpin is important for RNA binding (Fig. 5, Table 4, Suppl. Fig. S8) (Arthur et al. 2011; Holmqvist et al. 2018; Stein et al. 2020; Kim et al. 2022). Our studies showed that the minimal length of double-stranded stem of terminator in AniS-3ʹ and *rpmG*-3ʹ, which was sufficient for tight binding, was 2 G-C pairs, although in *rpmG*-3ʹ this minimal terminator stem was additionally extended by three A-U pairs including the uridines of the 3ʹ tail (Fig. 5A,B, Table 4, Suppl. Fig. S8A). The essential role of the 2 lowest G-C base pairs was also showed for the binding of truncated *malM*-3ʹ mutants to the FinO domain of *E. coli* ProQ (Stein et al. 2020). Additionally, the role of the lower part of the terminator in FinP RNA binding by FinO protein was shown using RNase footprinting (Arthur et al. 2011). The importance of the lower part of the terminator is also supported by the observation that disruption of three C-G base pairs, including a closing base pair of the terminator stem of the 3ʹ-UTR of *cspE* mRNA, abolished its binding by *S. enterica* ProQ (Holmqvist et al. 2018). Finally, the experiments with mutants of RocR RNA showed that shortening of its terminator stem to 5 base pairs did not weaken its binding to the FinO domain of *L. pneumophila* RocC. This observation was consistent with the crystal structure, which showed that several amino acid residues of the FinO domain of *L. pneumophila* RocC are within hydrogen bonding distance to the lowest 5 base pairs of the terminator of RocR (Kim et al. 2022).

There are subtle differences in the length of the 3ʹ tail, which is optimal for tight RNA binding by *N. meningitidis* ProQ and by other FinO domain proteins. For *rpmG*-3ʹ the constructs with the tail length of five to nine nucleotides bound tightly to *N. meningitidis* ProQ, but shortening the tail below five uridines gradually weakened the binding (Fig. 3A,B, Table 2, Suppl. Fig. S6A). Additionally, extending the length of the 3ʹ tail of *rpmG*-3ʹ to 13 residues essentially abolished the binding (Suppl. Fig. S3). In contrast, for AniS-3ʹ the tightest binding was observed for the 3ʹ tail length of six or seven uridines, while elongating the tail to eight uridines weakened the binding, and shortening the tail to less than four uridines abolished the binding (Fig. 3C,D, Table 2, Suppl. Fig. S6B). The presence of U-rich 3ʹ tails in the binding sites of *E. coli* ProQ and *S. enterica* ProQ was previously detected by CLIP-seq and RIL-seq studies (Holmqvist et al. 2018; Melamed et al. 2020). It was also shown that shortening of the 3ʹ tails of *malM*-3ʹ and *cspE*-3ʹ RNAs weakened their binding by *E. coli* ProQ (Stein et al. 2020). However, there were differences in the minimal tail lengths sufficient for *E. coli* ProQ binding by different RNAs, because for *malM*-3ʹ the 3ʹ tail length of 4 single-stranded uridines was sufficient for tight binding, while for *cspE*-3ʹ the 3ʹ tail length of 6 uridines, which included 2 single-stranded and 4 double stranded residues, was necessary for tight binding (Stein et al. 2020). The essential importance of the 3ʹ tail was also shown for the binding of FinP RNA by the FinO protein, where truncating the 3ʹ tail of FinP from GAU_4_ to only GA essentially abolished the binding (Jerome and Frost 1999). Additionally, it was also observed using gelshift assay that the binding affinities of FinO protein to RNAs derived from RocR, which had had the 3ʹ tail lengths of 3, 5, or 8 uridines, were quite similar (Kim et al. 2022). In contrast, the optimal tail length of RocR for binding to *L. pneumophila* RocC FinO domain was 5 nucleotides, and either shortening or elongating it strongly decreased the binding (Kim et al. 2022). This preference for a specific length of the 3ʹ tail was explained by the crystal structure which showed that the two terminal residues of the 3ʹ tail of RocR form hydrogen bonds with conserved residues of the FinO domain of RocC (Kim et al. 2022). The fact that the terminal residue of the 3ʹ tail has to be appropriately positioned for binding to conserved residues in the binding pocket of the FinO domain may be an important factor determining the optimal length of the tail, because either too short or too long 3ʹ tails would not be correctly positioned for these interactions. This suggests that the differences observed in the length of the 3ʹ tail optimal for tight RNA binding to different FinO-domain proteins could result from differences in the RNA binding sites in different FinO domains or from differences in RNA sequence or structure, which could affect the positioning of the terminus of the 3ʹ tail in relation to the binding pocket in the FinO domains.

Our data showed that modifications of 2ʹ-OH and 3ʹ-OH groups of the 3ʹ terminal ribose are strongly detrimental for the binding of RNAs derived from *rpmG*-3ʹ and AniS-3ʹ to *N. meningitidis* ProQ (Fig. 6, Table 5, Suppl. Fig. S9). The removal of both 2ʹ-OH and 3ʹ-OH groups abolished the ProQ binding of *rpmG*-ddC and AniS-ddC mutants. The binding was also weakened by the removal of the 3ʹ-OH group only or by blocking of the 2ʹ-OH group by a phosphate (Fig. 6, Table 5, Suppl. Fig. S9). The 2ʹ-OH and 3ʹ-OH groups of the terminal nucleoside are also important for RNA binding by other FinO-domain proteins. The gelshift-monitored RNA binding to the F-like plasmid FinO protein was abolished by the phosphorylation of the 3ʹ-OH group, and strongly weakened by the blocking of both hydroxyl groups by 2ʹ,3ʹ-dialdehyde (Arthur et al. 2011). The phosphorylation of the 3ʹ-OH group also abolished RocR RNA binding by *L. pneumophila* RocC, which was monitored using isothermal titration calorimetry (Kim et al. 2022). The reason for the importance of these hydroxyl groups in the binding has been explained by the crystal structure of *L. pneumophila* RocC, which showed that the 2ʹ-OH and 3ʹ-OH groups of the terminal uridine are within hydrogen bonding distance to peptide bond amino groups of conserved amino acid residues in type II β-turn between helices 3 and 4 of RocC (Kim et al. 2022). Interestingly, the contribution of the 3ʹ-terminal 3ʹ-OH group to RNA binding has been also observed for *Salmonella enterica* Hfq (Sauer and Weichenrieder 2011), which suggests that contacts with the hydroxyl groups of the 3ʹ terminal nucleoside are a common feature of bacterial proteins which recognize RNAs at their 3ʹ ends.

The substitution of an A-rich stretch on the 5ʹ side of the terminator of *rpmG*-3ʹ weakens its binding to *N. meningitidis* ProQ, while the substitution of an U-rich stretch in corresponding position of AniS-3ʹ strengthens its binding to ProQ (Fig. 7, Table 6, Suppl. Fig. S10). We have previously observed that in RNA ligands of ProQ identified in *E. coli* and *S. enterica* the sequence of the corresponding region is enriched in adenosine residues, as opposed to RNA ligands of Hfq, where this region is uridine-enriched (Holmqvist et al. 2018; Melamed et al. 2020; Stein et al. 2020). However, when we compared the corresponding sequence in RNAs bound by *N. meningitidis* ProQ (Bauriedl et al. 2020) and Hfq (Heidrich et al. 2017), we found that in both of these groups of RNAs this sequence is A-enriched, and that the adenosine enrichment of this region is a general feature of *N. meningitidis* transcriptome (Suppl. Fig. S5, Suppl. Table S4). We note, that because the identification of Hfq ligands in *N. meningitidis* was obtained using RIP-seq (Heidrich et al. 2017), it is not possible to distinguish whether the Hfq binding sites were located at the terminator structures or elsewhere in the sequence. Interestingly, among 40 RNAs bound by Hfq, which were included in our sequence logo analysis, are sRNAs AniS, RcoF1, RcoF2, and 3ʹ-UTR of NMV_1651, which have U-rich motifs in this region (Suppl. Table S4) (Fantappie et al. 2011; Heidrich et al. 2017; Bauriedl et al. 2020). The fact that substitution of uridines to adenosines in the region improved the ProQ binding to AniS, while the substitution of adenosines to uridines in the corresponding region weakened the binding of *rpmG*-3ʹ supports the importance of the sequence 5ʹ adjacent to the terminator to RNA recognition by *N. meningitidis* ProQ.

Overall, our studies showed that the minimal ProQ from *N. meningitidis* recognizes RNAs in a generally similar way as the isolated FinO domains from other FinO-domain proteins. However, there are certain differences between them, which are related mainly to the sequence and length of single-stranded RNA sequences surrounding the terminator hairpin, which are required for optimal binding by *N. meningitidis* ProQ.

## MATERIALS AND METHODS

### Protein preparation

The sequence of *N. meningitidis* ProQ was cloned from pTYB11-ProQMenningo (a kind gift of Prof. Jörg Vogel, University of Würzburg) into expression vector pET-15b by PCR amplification and restriction digestion with BamH1 in *E. coli* DH5α. The resulting construct had N-terminal cleavable His_6_-tag followed by TEV protease recognition site. After cleavage ProQ had a single additional serine residue on the N-terminus. The expression plasmid was transformed into *E. coli* BL21 Δ*hfq* strain (a kind gift of Prof. Agnieszka Szalewska-Pałasz, University of Gdańsk). *N. meningitidis* ProQ was purified essentially as described previously for *E. coli* ProQ (Stein et al. 2020). In short, N-terminally His_6_-tagged ProQ was purified using nickel affinity chromatography, which was followed by heparin affinity chromatography to remove contaminating nucleic acids. After cleaving off the His_6_-tag using TEV protease, the tag was removed using the second nickel affinity chromatography, which was followed by size-exclusion chromatography. The purified ProQ was stored in buffer consisting of 50 mM Tris, pH 7.5, 300 mM NaCl, 10% glycerol, and 1 mM EDTA, in 10 µl aliquots of 10 μM concentration at -80 °C. The aliquots were used without refreezing. The molecular weight of the purified protein with additional N-terminal serine residue remaining from TEV cleavage site was determined by MALDI-TOF as 15614.9 Da, which agrees with the calculated mass of 15614.7 Da. The protein concentration was determined by measuring the absorption at 280 nm using extinction coefficient of 5240 M^−1^ cm^−1^.

### RNA preparation

The DNA templates used for *in vitro* transcription were obtained by Taq polymerase extension of chemically synthesized overlapping oligodeoxyribonucleotides (Metabion) (Supplemental Table S2). RNA molecules were transcribed with T7 RNA polymerase and purified using denaturing gel electrophoresis, as described (Milligan et al. 1987; Olejniczak 2011). In the next step, RNAs were 5′-^32^P labeled using T4 polynucleotide kinase (Thermo Scientific), followed by phenol-chloroform extraction, purification using denaturing gel and precipitation with ethanol. The obtained RNAs were dissolved in water and stored at -20 °C.

Chemically synthesized RNA oligos (Metabion) were purified with denaturing gel electrophoresis followed by ^32^P-labeling (Supplemental Table S3).

### RNA binding assay

Before use, RNA molecules were denatured for 2 min at 90°C followed by 5 min refolding on ice. The concentration series of *N. meningitidis* ProQ was prepared by two-fold dilutions from the concentration of 50 nM. In all binding reactions, 1 nM ^32^P-labeled RNA was mixed with the protein sample in binding buffer consisting of 25 mM Tris, pH 7.5, 150 mM NaCl, 5% glycerol, and 1 mM MgCl_2_, and incubated for 30 min at room temperature in low-protein binding microplates pretreated with 0.0025% bovine serum albumin solution. After incubation, reactions were loaded onto a 6% polyacrylamide gel (19:1) at 4°C. After the electrophoresis, gels were vacuum-dried, and exposed to phosphor screens overnight. The signal was quantified using a phosphorimager (Fuji FLA-5000) and MultiGauge software, and data were fitted to a quadratic equation using GraphPad Prism software. The average *K*_d_ values were calculated from at least three independent experiments.

## ACKNOWLEDGEMENTS

We thank Gisela Storz for helpful discussions, and Julia Kurzawska, Maria Mamońska, Monika Mazur and Joanna Zwolenkiewicz for critical comments on the manuscript. This work was supported by National Science Centre in Poland [grants No. 2021/41/N/NZ1/04133 to M.M.B., and No. 2018/31/B/NZ1/02612 and No. 2022/47/B/NZ1/01665 to M.O.]. Funding for open access charge: National Science Centre [2022/47/B/NZ1/01665] and Adam Mickiewicz University.

## AUTHORS CONTRIBUTIONS

M.M.B. performed all experiments, M.M.B. and M.O. analyzed the data and wrote the manuscript.

