## Supplemental Materials for "RNA recognition by minimal ProQ from *Neisseria meningitidis*"

### SUPPLEMENTAL MATERIALS AND METHODS

#### ProQ-binding assay

The RNA binding to ProQ protein was measured using a gelshift assay as described in the Materials and Methods section in the main text.

#### Analysis of RNA sequences

To compare the nucleotide composition of the sequence 5′-adjacent to the terminator stem, we used data from global profiling studies using CLIP-seq for *N. meningitidis* ProQ (Bauriedl et al. 2020) and using RIP-seq for *N. meningitidis* Hfq (Heidrich et al. 2017). For the analysis, we selected 40 3′-UTRs and sRNAs, which were the top ligands containing Rho-independent terminators of each protein (Suppl. Fig. S4). In CLIP-seq data for ProQ-bound RNAs only those RNAs were selected, in which the peak at least partially covered a terminator hairpin. In RIP-seq data for Hfq-bound RNAs all RNAs annotated as 3' UTRs or sRNAs were considered in the analysis. In these RNA ligands, a 10-nt long sequence 5′-adjacent to the intrinsic terminator was analyzed. To define the sequences 5'-adjacent to the terminator hairpins the secondary structure of each RNA was analyzed using the *ViennaRNA* package (Lorenz et al. 2011). The 3' end of this sequence was defined as the nucleotide located immediately on the 5' side of the closing G-C/C-G pair of the terminator stem. The extracted 10-nt long sequences were then used to generate the sequence logos with *WebLogo* (Crooks et al. 2004). RNA sequences selected for analysis are shown in Supplemental Table S4.

Analogous analysis was carried out for a random gene sample of 98 genes from the *Neisseria meningitidis* transcriptome. Subsequently, the statistical importance of nucleotide frequency at each position of the 10-nt sequence 5'-adjacent to the terminator hairpin in the RNA ligands of ProQ was calculated in reference to the RNA ligands of Hfq and the random gene sample using the *pLogo* software (O'Shea et al. 2013).

### SUPPLEMENTAL TABLES

**Supplemental Table S1. DNA oligonucleotides used to prepare the insert to clone *Neisseria meningitidis proQ* gene into the pET-15b expression plasmid.**

| name | sequence (5' → 3') |
|---|---|
| Nm-ProQ-F | cggc**ggatcc**g**gaaaatctttatttccaatcc**ATGACACAAGAAACCGCTTTGGGTGC |
| Nm-ProQ-R | cggc**ggatcc**TTATTCTGCTGCGGAAGATTCGGCTGC |

cggc - sequence inserted to improve the efficiency of BamHI cleavage

**ggatcc -** BamHI restriction site

**gaaaatctttatttccaatcc -** sequence encoding amino acid sequence recognised by protease TEV (ENLYFQ↓S)

**Supplemental Table S2. Oligonucleotides used to prepare templates for *in vitro* transcription in primer extension reaction.**

| name | sequence (5' → 3') |
|---|---|
| AniS-2UtoA-F | **TAATACGACTCACTATA**GGTTTCCTCTTGATGTGTGTGTGTAAGGGTGTGG |
| AniS-2U-R | AAGGGGTGGCGGCAGCCACACCCAAACACACACAC |
| AniS-2UtoA-R | AAAAAAAAGGGGTGGCGGCAGCCACACCCTTACACACACAC |
| AniS-3UtoA-F | **TAATACGACTCACTATA**GGTTTCCTCTTGATGTGTGTGTGAAAGGGTGTGGCTGC |
| AniS-3UtoA-R | AAAAAAAAGGGGTGGCGGCAGCCACACCCTTTCACAC |
| AniS-3'-F | **TAATACGACTCACTATA**GTTTCCTCTTGATGTGTGTGTGTTTGGGTGTGG |
| AniS-3'-R | AAAAAAAAGGGGTGGCGGCAGCCACACCCAAACACACACAC |
| AniS-noU-R | GGGGTGGCGGCAGCCACACCCAAACACACACAC |
| AniS-U2-R | AAGGGGTGGCGGCAGCCACACCCAAACACACACAC |
| AniS-U4-R | AAAAGGGGTGGCGGCAGCCACACCCAAACACACACAC |
| AniS-U5-R | AAAAAGGGGTGGCGGCAGCCACACCCAAACACACACAC |
| AniS-U6-R | AAAAAAGGGGTGGCGGCAGCCACACCCAAACACACACAC |
| AniS-U7-R | AAAAAAAGGGGTGGCGGCAGCCACACCCAAACACACACAC |
| Bns1-F | **TAATACGACTCACTATA**GGACAAGTCCAAGTATTATAAAGGCTGAATAAAGAGGAAACAGCAGGCAGATATA |
| Bns1-R | AAAAAAGCAGATATATTCGGACTGCACCTCCCGAATATATCTGCCTGCTGTTTCC |
| *carA*-5'-F | **TAATACGACTCACTATA**GGGTATAGGCACTTGCCCGAAAAGCACGTTACG |
| *carA*-5'-R | AAAAAAAAACACGCCGCGCAAGATAGACGCGTAACGTGCTTTTCGGGCA |
| *iga*-3'-F | **TAATACGACTCACTATA**GAAAATACTAAATTCATAGCAAAATAAAATGCCGTCTGA |
| *iga*-3'-R | ATAAAAATGCCGTCTGAAGCCTGAGTTCAGACGGCATTTTATTTTG |
| *intergenic*-F | **TAATACGACTCACTATA**GGATATGCCGAAAGGGGTTTGACGATGCCGCCGTGCGCTGTC |
| *intergenic*-R | AGAAAAAGGGTTTGACAGCGCACGGCGGCATCG |
| *pnp*-3'ext-F | **TAATACGACTCACTATA**GTTTACAACCTGCCGCACGGTCTGAAACCCTAACTATGCACATTCGGATTTTA |
| *pnp*-3'ext-R | TATTGTTCCTTTCAAAATACCGCACTGCTAAAACACTAATAATGCACACTAAAATCCGAATGTGCATAG |
| *pnp*-5'-F | **TAATACGACTCACTATA**GTTTACAACCTGCCGCACGGTCTGAAACCCTAACTATG |
| *pnp*-5'-R | AATAATGCACACTAAAATCCGAATGTGCATAGTTAGGGTTTCAGACC |
| *rpmG*-2AtoU-F | **TAATACGACTCACTATA**GGGATTTCAGTTTGAAAGCATTGCCTCCGA |
| *rpmG*-2AtoU-R | AAAAAAGCCTCCGAACAGTCGGAGGCAATGCTTTCAAA |
| *rpmG*-31-F | **TAATACGACTCACTATA**GCAAAGCCTCCGACTGTTC |
| *rpmG*-31-R | AAAAAAGCCTCCGAACAGTCGGAGGCTTTGC |
| *rpmG*-33-F | **TAATACGACTCACTATA**GAGCAAAGCCTCCGACTGTTC |

| | |
|---|---|
| *rpmG*-35/33-R | AAAAAAGCCTCCGAACAGTCGGAGGCTTTGCTT |
| *rpmG*-35-F | **TAATACGACTCACTATA**GAAAGCAAAGCCTCCGACTGTTC |
| *rpmG*-36-loop-F | **TAATACGACTCACTATA**GGGATTTCAGTTTGAAAGCAAAGCGAAAG |
| *rpmG*-36-loop-R | AAAAAAGCTTTCGCTTTGCTTTCAAAC |
| *rpmG*-3AtoU-F | **TAATACGACTCACTATA**GGGATTTCAGTTTGAAAGCTTTGCCTCCGA |
| *rpmG*-3AtoU-R | AAAAAAGCCTCCGAACAGTCGGAGGCAAAGCTTTCAAA |
| *rpmG*-3UtoA-R | AAATAAGCCTCCGAACAGTCGGAGGCTTTGC |
| *rpmG*-3'ext-R | ATAAAAATAAAAAAGCCTCCGAACAGTCGGAGGCTTTGCTTTCAAA |
| *rpmG*-3'-F | **TAATACGACTCACTATA**GGGATTTCAGTTTGAAAGCAAAGCCTCCGA |
| *rpmG*-3'-R | AAAAAAGCCTCCGAACAGTCGGAGGCTTTGCTTTCAAA |
| *rpmG*-40-loop-F | **TAATACGACTCACTATA**GGGATTTCAGTTTGAAAGCAAAGCCTGAAAAGG |
| *rpmG*-40-loop-R | AAAAAAGCCTTTTCAGGCTTTGCTTTC |
| *rpmG*-44-loop-F | **TAATACGACTCACTATA**GGGATTTCAGTTTGAAAGCAAAGCCTC |
| *rpmG*-44-loop-R | AAAAAAGCCTCCTTTCGGAGGCTTTGCTTTCAAACTG |
| *rpmG*-7U6A-R | TAAAAAAGCCTCCGAACAGTCGGAGGCTTTGCTTTCAAA |
| *rpmG*-95-F | **TAATACGACTCACTATA**GATCCGGTTGCCCGCAAACACGTAGTGTACAAAGAAACCAAACTGAAATAATTT |
| *rpmG*-F | **TAATACGACTCACTATA**GGGATTTCAGTTTGAAAGCAAAGCCTCCGA |
| *rpmG*-loop-F | **TAATACGACTCACTATA**GGGAUUUCAGUUUGAAAGCAAAGCCUCCGAGAA |
| *rpmG*-loop-R | GAAAGCAAAGCCUCCGAGAAAUCGGAGGCUUUUUU |
| *rpmG*-noU-R | GCCTCCGAACAGTCGGAGGCTTTGCTTTCAAA |
| *rpmG*-R | AAAAAAGCCTCCGAACAGTCGGAGGCTTTGCTTTCAAA |
| *rpmG*-U2 | AAGCCTCCGAACAGTCGGAGGCTTTGCTTTCAAA |
| *rpmG*-U4-R | AAAAGCCTCCGAACAGTCGGAGGCTTTGCTTTCAAA |
| *rpmG*-U5-R | AAAAAGCCTCCGAACAGTCGGAGGCTTTGCTTTCAAA |
| *rpmG*-U6AU2-R | AATAAAAAAGCCTCCGAACAGTCGGAGGCTTTGCTTTCAAA |
| *rpmG*-U6AU-R | ATAAAAAAGCCTCCGAACAGTCGGAGGCTTTGCTTTCAAA |
| *rpmG*-UtoA-F | **TAATACGACTCACTATA**GGGATTTCAGTTTGAAAGCAAAGCCTCCGACTG |
| *sodC*-3'ext-R | ATATCTAGCAAAAAAGTGCGGTCAAAATTCACCGCACTTTTCGTTTCGAACA |
| *sodC*-3'-F | **TAATACGACTCACTATA**GATTAAATAATTCGATTGTTCGAAACGAAAAGTGCG |
| *sodC*-3'-R | AAAAAAGTGCGGTCAAAATTCACCGCACTTTTCGTTTCGAACA |

The **bold font** is the sequence of T7 polymerase promoter.

**Supplemental Table S3. Oligoribonucleotides obtained by chemical synthesis (Metabion)**

| **name** | **sequence (5' → 3')** |
|---|---|
| AniS-23 | UGUGUGUGUUUGAAAUUUUUUUU |
| AniS-25 | UGUGUGUGUUUGGAAACUUUUUUUU |
| AniS-29 | GGGUGUGGCUGCCGCCACCCCUUUUUUUU |
| AniS-31 | UUGGGUGUGGCUGCCGCCACCCCUUUUUUUU |
| AniS-33 | GUUUGGGUGUGGCUGCCGCCACCCCUUUUUUUU |
| AniS-35 | GUGUUUGGGUGUGGCUGCCGCCACCCCUUUUUUUU |
| AniS-37 | GUGUGUUUGGGUGUGGCUGCCGCCACCCCUUUUUUUU |
| AniS-40 | UGUGUGUGUUUGGGUGUGGCUGCCGCCACCCCUUUUUUUU |
| AniS-loop | UGUGUGUGUUUGGGUGUGGCGAAAGCCACCCCUUUUUUUU |
| AniS-loop-27 | UGUGUGUGUUUGGGAAACCUUUUUUUU |
| AniS-loop-29 | UGUGUGUGUUUGGGGAAACCCUUUUUUUU |
| AniS-loop-31 | UGUGUGUGUUUGGGGGAAACCCCUUUUUUUU |
| AniS-loop-35 | UGUGUGUGUUUGGGGUGGAAACACCCCUUUUUUUU |
| AniS-loop-39 | UGUGUGUGUUUGGGGUGGCGAAAGCCACCCCUUUUUUUU |
| *rpmG*-26 | GCCUCCGACUGUUCGGAGGCUUUUUU |
| *rpmG*-29 | AAAGCCUCCGACUGUUCGGAGGCUUUUUU |
| *rpmG*-45 | AUUUCAGUUUGAAAGCAAAGCCUCCGACUGUUCGGAGGCUUUUUU |

**Supplemental Table S4. 10 nt sequences on the 5' side of the closing G-C/C-G pair of terminator stems of ProQ- and Hfq-specific RNAs, which were used for the sequence logo analysis.**

| Top 40 *N. meningitidis* ProQ ligands CLIP-seq dataset (Bauriedl et al. 2020) | sequence | Top 40 *N. meningitidis* Hfq ligands RIP-seq dataset (Heidrich et al. 2017) | sequence |
|---|---|---|---|
| intergenic, 2094444-2094518(-) | AUGCCGAAAG | NMV_1189 3' | UUGAAACAAU |
| iga 3' | AAAAUAAAAU | recC 3' | AAAGGCAAAU |
| NMV_1592 3' | AUAAGGCAAU | uvrB 3' | GAGUAAUAAA |
| comE1 3' | CGCAUCAAAU | recR 3' | AAUGCAAAAU |
| NMnc0006 | ACUACACAAA | NMV_1052 3' | UCUGGAAAAU |
| nuoN 3' | GCCGCCGCAA | prmB 3' | AAGGGGAAAU |
| NMV_1266 3' | AUCCGAACAA | RcoF1 | UGUAGACUUG |
| nrdB 3' | UUGAAAAAAU | RcoF2 | UGUAGACUUG |
| pilX 3' | UUUUGCCAAU | Int 3' | AGUGAUACCU |
| alx 3' | AAACAAAAAA | lpxA 3' | GCUGACCGUA |
| NMV_0646 3' | GUUAUGAAAA | NMV_1503 3' | UUUGUAUUAU |
| nagZ 3' | CAUAUUCAAA | NMV_1295 3' | CGAUUCUAAG |
| NMV_0508 3' | CAUUAAAAAU | NMV_0065 3' | UGAAAAAAAA |
| panC 3' | CGUCGGGAAU | NMV_2316 3' | AGCUAGGUAA |
| queA 3' | GAAGACAAGU | NMV_1651 3' | GUUCCUUUUG |
| intergenic, 765600-765668(-) | AAUAUCAAAG | NMV_2146 3' | UAUCUGCAAA |
| intergenic, 1916777-1916862(-) | GUUAAAAAAU | NMV_0065 3' | GAAGACAAGU |
| NMnc0041 | UUGGGGGGAA | NMnc0041 | UUGGGGGGAA |
| dnaN 3' | UGGAGAUAUA | NMV_0057 3' | CGCAUCAAAU |
| gloA 3' | CGCCGCCAAU | NMV_0638 3' | UUGAACGAAU |
| rplI 3' | AUUAUCAGAU | NMV_0320 3' | UAAUUUAAAA |
| NMV_0480 3' | GCAUCAAAAU | NMV_1804 3' | AAGCGCAUAU |
| NMV_0949 3' | UCGCAAAAAU | sigEsRNA | UCCUUGGGAA |
| purM 3' | AUAGCGAAAU | AniS | UGUGUGUUUU |
| NMV_2143 3' | CAUUGAAAAU | NMV_1296 3' | AGAAUGGAAU |
| carA 3' | GCCCGAAAAG | NMV_0692 3' | UAAGUCAAAA |
| dsbA2 3' | AAUGAGUAAA | NMV_0091 3' | UGACGGAAAU |
| NMV_0786 3' | AGCCCGAUAU | ppsA 3' | CAAUGCCCAU |

| | | | |
|---|---|---|---|
| intergenic, 1193850-1193917(+) | CAAAACAUAA | NMV_0441 3' | UUGACAGAAU |
| rplY 3' | GUUGUUUUAU | NMV_1746 3' | UUUGAUUAAA |
| dsbD 3' | UGUUUCAAAU | NMV_1201 3' | CGAUAAAAAA |
| atpC 3' | GUUUGGAAAA | rplS 3' | UUCCCCCAAU |
| intergenic, 2203594-2203659(-) | CCCAGAUAAA | NMV_1560 3' | AACGGCAAAU |
| NMV_0791 3' | GGAUUGAAAU | NMV_1509 3' | UUUUGCCAAU |
| NMnc0044 | GAAGACAAGU | NMV_2134 3' | CGGAAAAGGU |
| NMV_1640 3' | GUAAAAAAAU | NMV_0716 3' | AACCAUAAAG |
| dnaJ 3' | CGGAAACAAG | recA 3' | UCCCGAUAAA |
| prc 3' | UUUUUACGAU | dnaK 3' | UCUGAAAAAA |
| comP 3' | UUCGCCACUU | NMV_1336 3' | UUUUUGAAAU |
| intergenic, 2167518-2167594(+) | AAAUAUCAAA | bfr 3' | CAUCAUCAGA |

### SUPPLEMENTAL FIGURES

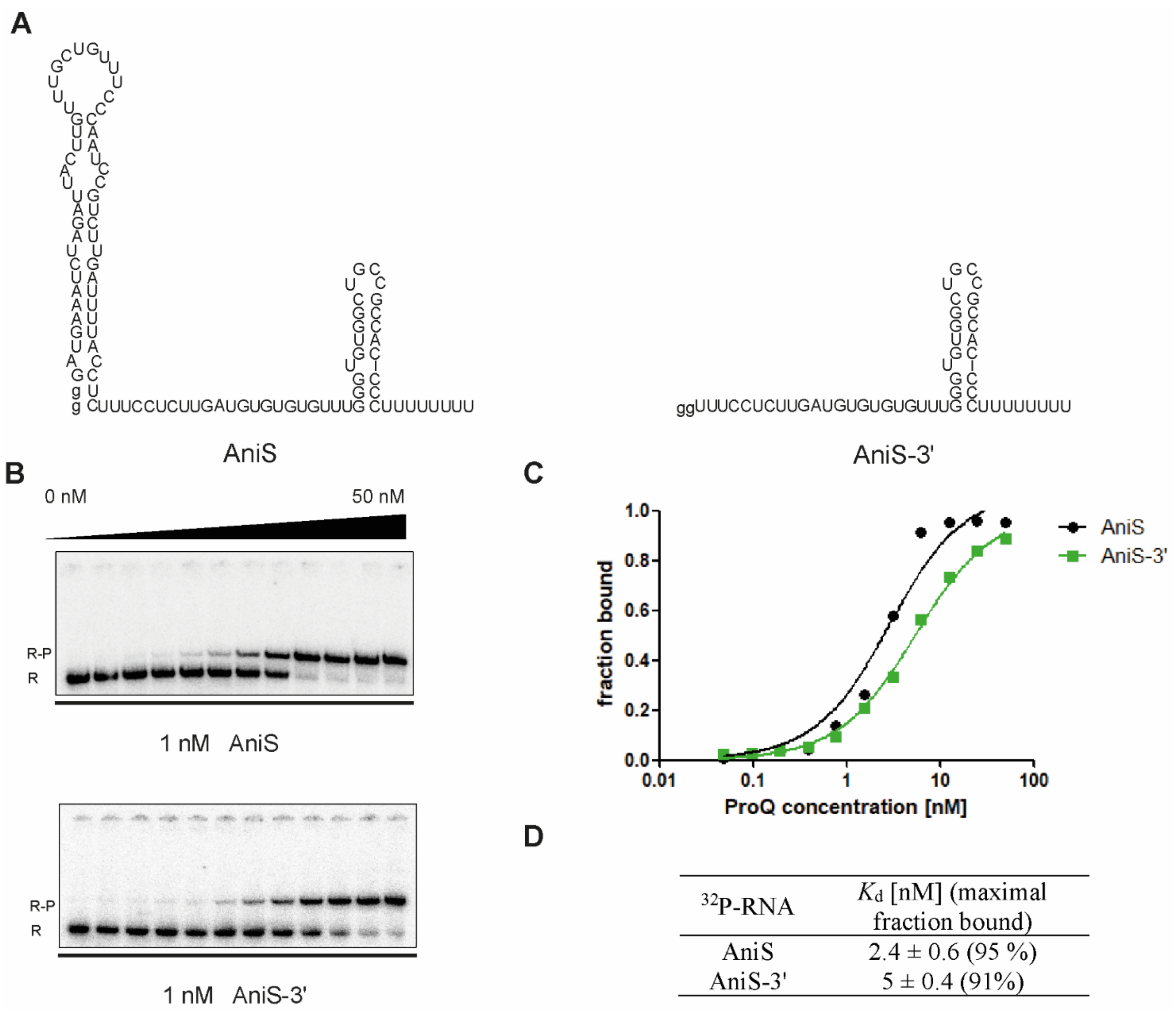


| $^{32}$P-RNA | $K_d$ [nM] (maximal fraction bound) |
|---|---|
| AniS | 2.4 ± 0.6 (95 %) |
| AniS-3' | 5 ± 0.4 (91%) |

**Supplemental Figure S1. The full-length AniS and its 3′-terminal fragment, named AniS-3′, have similar binding affinities to *N. meningitidis* ProQ**. (A) Secondary structures of full-length AniS and AniS-3'. (B) The binding of $^{32}$P-labeled RNAs AniS-3' and AniS was monitored using a gelshift assay. Free $^{32}$P-RNA is marked as R, RNA-ProQ complexes as R-P. (C) The fitting of the ProQ binding data from B using the quadratic equation. (D) The average equilibrium dissociation constant ($K_d$) values and the maximum RNA fraction bound calculated from at least three independent experiments. The ProQ binding data for AniS-3′ in (C) and (D) are the same as in Figure 2 and Table 1, respectively. The RNA secondary structure predictions were performed in the *ViennaRNA* program (Lorenz et al. 2011).

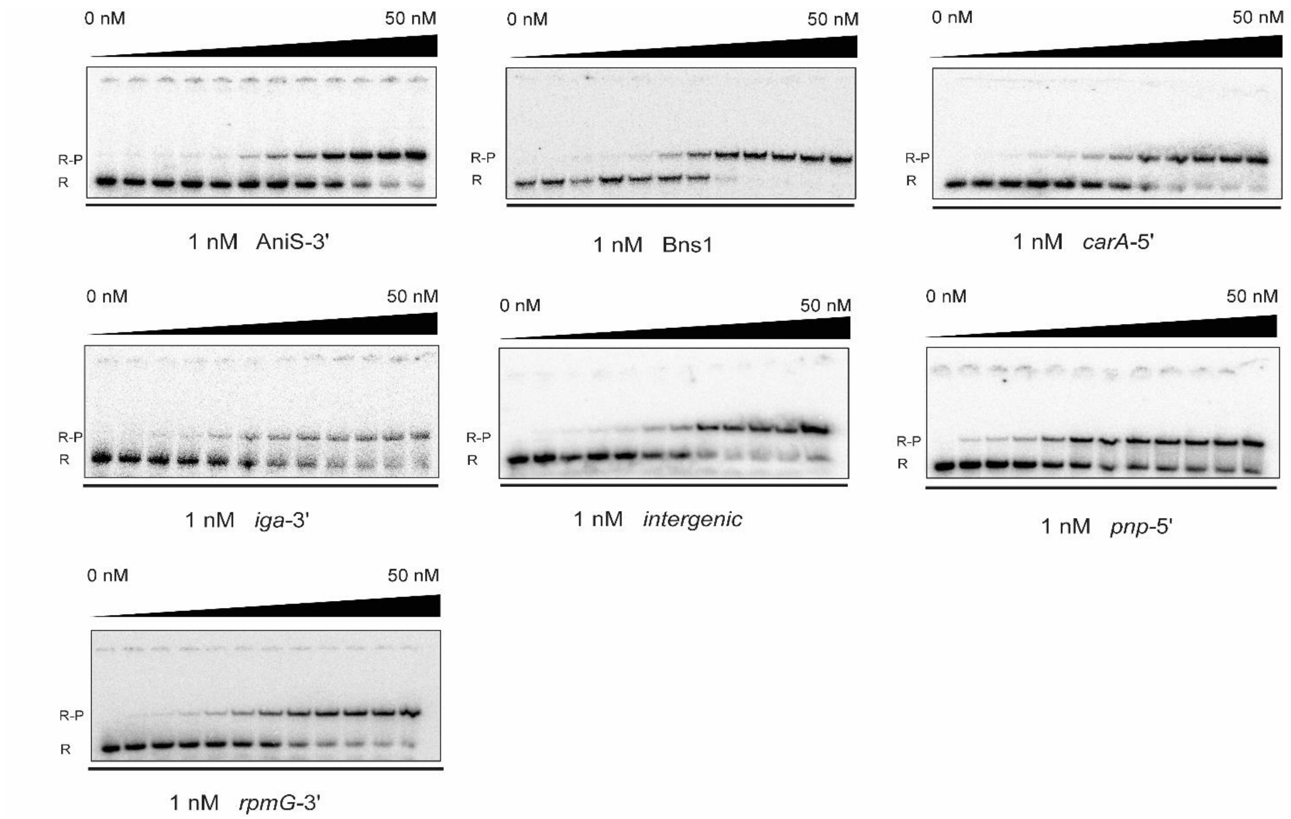


**Supplemental Figure S2. Gelshift analysis of the binding of $^{32}$P-labeled RNAs AniS-3', Bns1, *carA*-5', *iga*-3', *intergenic*, *pnp*-5' and *rpmG*-3' to *N. meningitidis* ProQ**. The raw data in the gels correspond to the data presented in plots in Fig. 2 in the main text. The concentration series of ProQ protein was made by 2-fold sequential dilutions. The range of the ProQ concentrations is indicated above the gels. Free $^{32}$P-RNA is marked as R, RNA-ProQ complex as R-P.

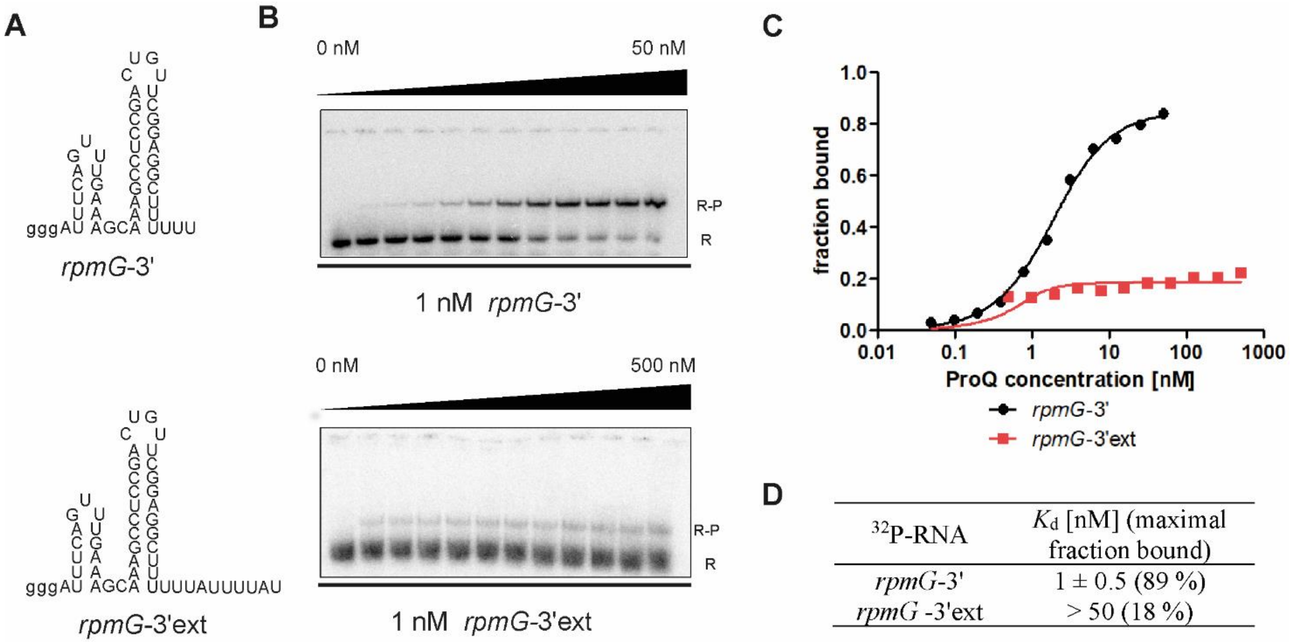


**Supplemental Figure S3. The 3′ extension of *rmpG*-3′ weakens its binding by *N. meningitidis* ProQ.** (A) Secondary structures of *rpmG*-3' and of *rpmG*-3'ext. (B) The binding of $^{32}$P-labeled RNAs *rpmG*-3' and *rpmG*-3'ext was monitored using a gelshift assay. Free $^{32}$P-RNA is marked as R, RNA-ProQ complexes as R-P. (C) The fitting of the ProQ binding data from B using the quadratic equation. (D) The average equilibrium dissociation constant ($K_d$) values and the maximum RNA fraction bound calculated from at least three independent experiments. The ProQ binding data for *rpmG*-3′ in (C) and (D) are the same as in Figure 2 and Table 1, respectively. The RNA secondary structure predictions were performed in the *ViennaRNA* program (Lorenz et al. 2011).

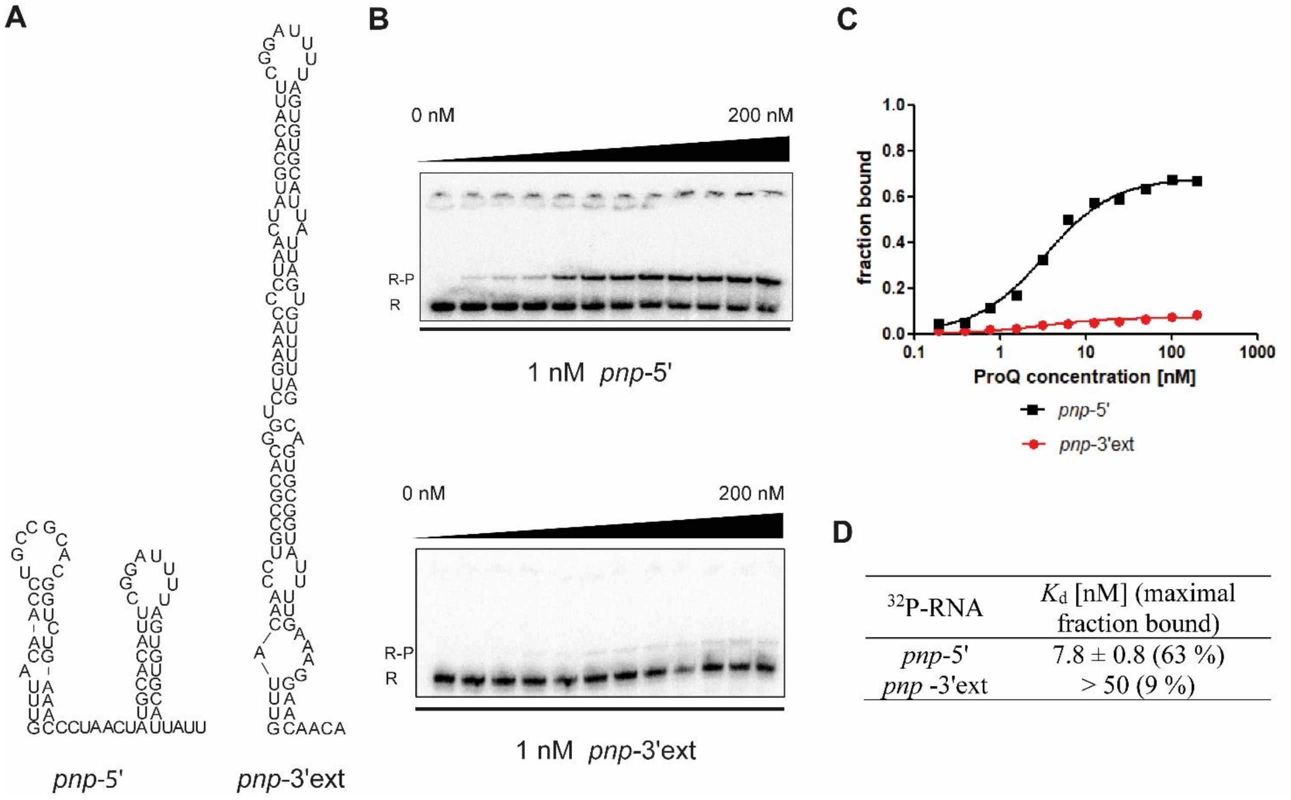


**Supplemental Figure S4. The 3′ extension of *pnp*-5' weakens its binding by *N. meningitidis* ProQ**. (A) Secondary structures of *pnp*-5' and *pnp*-3'ext. (B) The binding of $^{32}$P-labeled RNAs *pnp*-5' and *pnp*-3'ext was monitored using a gelshift assay. Free $^{32}$P-RNA is marked as R, RNA-ProQ complexes as R-P. (C) The fitting of the ProQ binding data from B using the quadratic equation. (D) The average equilibrium dissociation constant ($K_d$) values and the maximum RNA fraction bound calculated from three independent experiments. The average $K_d$ value for *pnp*-5′ is the same as in Table 1 in the main text. The RNA secondary structure predictions were performed in the *ViennaRNA* program (Lorenz et al. 2011).

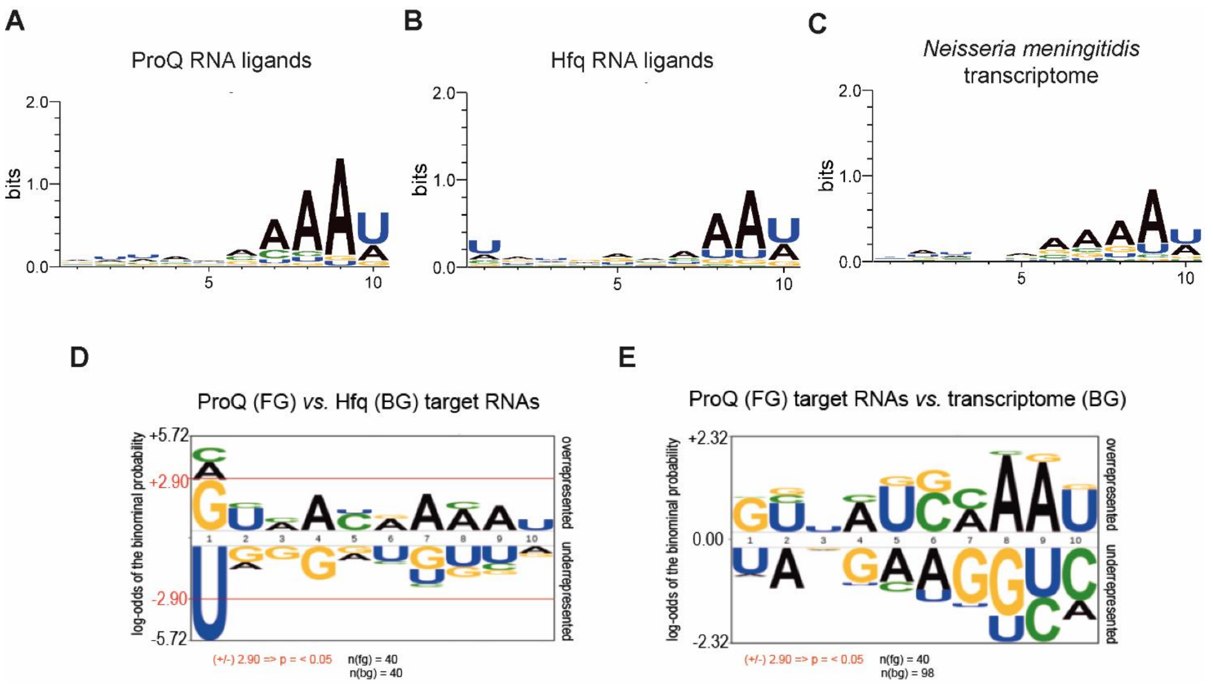


**Supplemental Figure S5. The comparison of the nucleotide compositions of the 10-nt sequence on the 5' side of the closing G-C/C-G pair of the intrinsic terminator stem in RNA ligands of ProQ and Hfq, and in a random gene sample of *N. meningitidis*.** Nucleotide frequencies in the 10-nt long sequence 5' adjacent to the terminator hairpin are shown for the top 40 RNA ligands of ProQ and Hfq containing Rho-independent terminators, in the data obtained previously by CLIP-seq for ProQ (Bauriedl et al. 2020) and RIP-seq for Hfq (Heidrich et al. 2017), in *N. meningitidis*. The last RNA ligand used for ProQ ligands logo generation (#40) corresponds to RNA positioned 74$^{th}$ in the CLIP-seq data set of 235 RNA ligands. The last RNA ligand used for Hfq ligands logo generation (#40) corresponds to RNA positioned 418$^{th}$ in the RIP-seq data set of 615 RNA ligands having logFC above 1.0. Nucleotide frequencies for *N. meningitidis* transcriptome were obtained for a random sample of 98 genes from *N. meningitidis*. (A) Nucleotide frequencies obtained by the *WebLogo* software for the top 40 *N. meningitidis* ProQ ligand RNAs containing Rho-independent terminators in CLIP-seq data. (B) Nucleotide frequencies obtained by the *WebLogo* software for the top 40 *N. meningitidis* Hfq ligand RNAs containing Rho-independent terminators in RIP-seq data. (C) Nucleotide frequencies obtained by *WebLogo* software for 98 randomly selected transcripts of *N. meningitidis*. The statistical significance of nucleotide frequencies at each nucleotide position for ProQ RNA ligands as compared to Hfq RNA ligands (D) and for ProQ RNA ligands as compared to the random sample of transcripts (E). Sequences were analysed by *pLogo* software for the top 40 *N. meningitidis* ProQ RNA ligands as the foreground, and the top 40 Hfq target RNAs or 98 gene sample from *N. meningitidis* genome as the backgrounds. Statistically significant P-value was only observed for U at position 1, P = 0.005 in ProQ ligands *vs.* Hfq ligands. The number of foreground sequences was marked on the figures as n(fg), and the number of background sequences as n(bg).

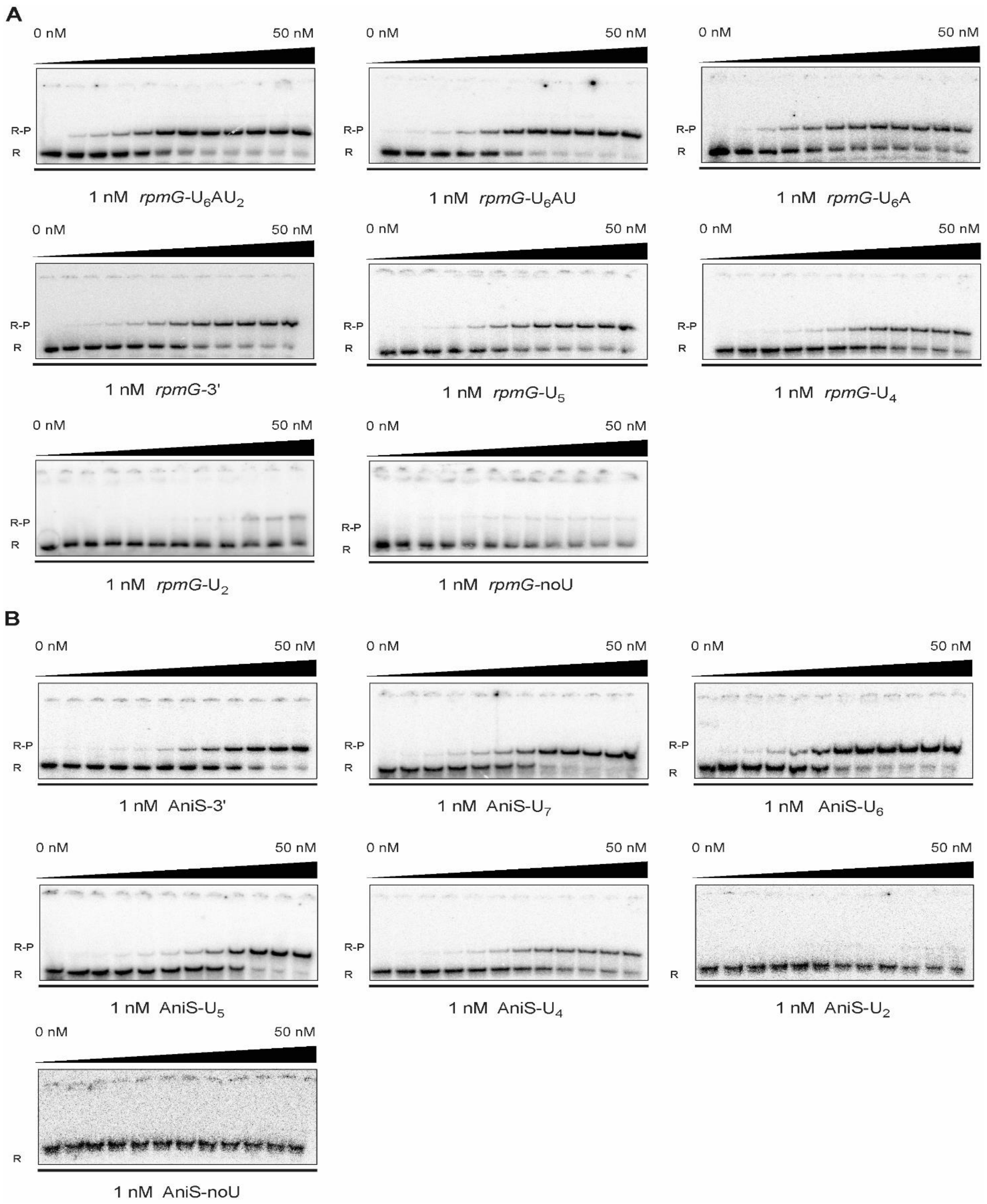


**Supplemental Figure S6. Gelshift analysis of the binding of $^{32}$P-labeled RNAs (A) *rpmG*-U$_6$AU$_2$, *rpmG*-U$_6$AU, *rpmG*-U$_6$A, *rpmG*-3', *rpmG*-U$_5$, *rpmG*-U$_4$, *rpmG*-U$_2$ and *rpmG*-noU, (B) AniS-3', AniS-U$_7$, AniS-U$_6$, AniS-U$_5$, AniS-U$_4$, AniS-U$_2$ and AniS-noU to *N. meningitidis* ProQ**. The raw data in the gels correspond to the data presented in plots in Fig. 3 in the main text. The concentration series of ProQ protein was made by 2-fold sequential dilutions. The range of the ProQ concentrations is indicated above the gels. Free $^{32}$P-RNA is marked as R, RNA-ProQ complex as R-P.

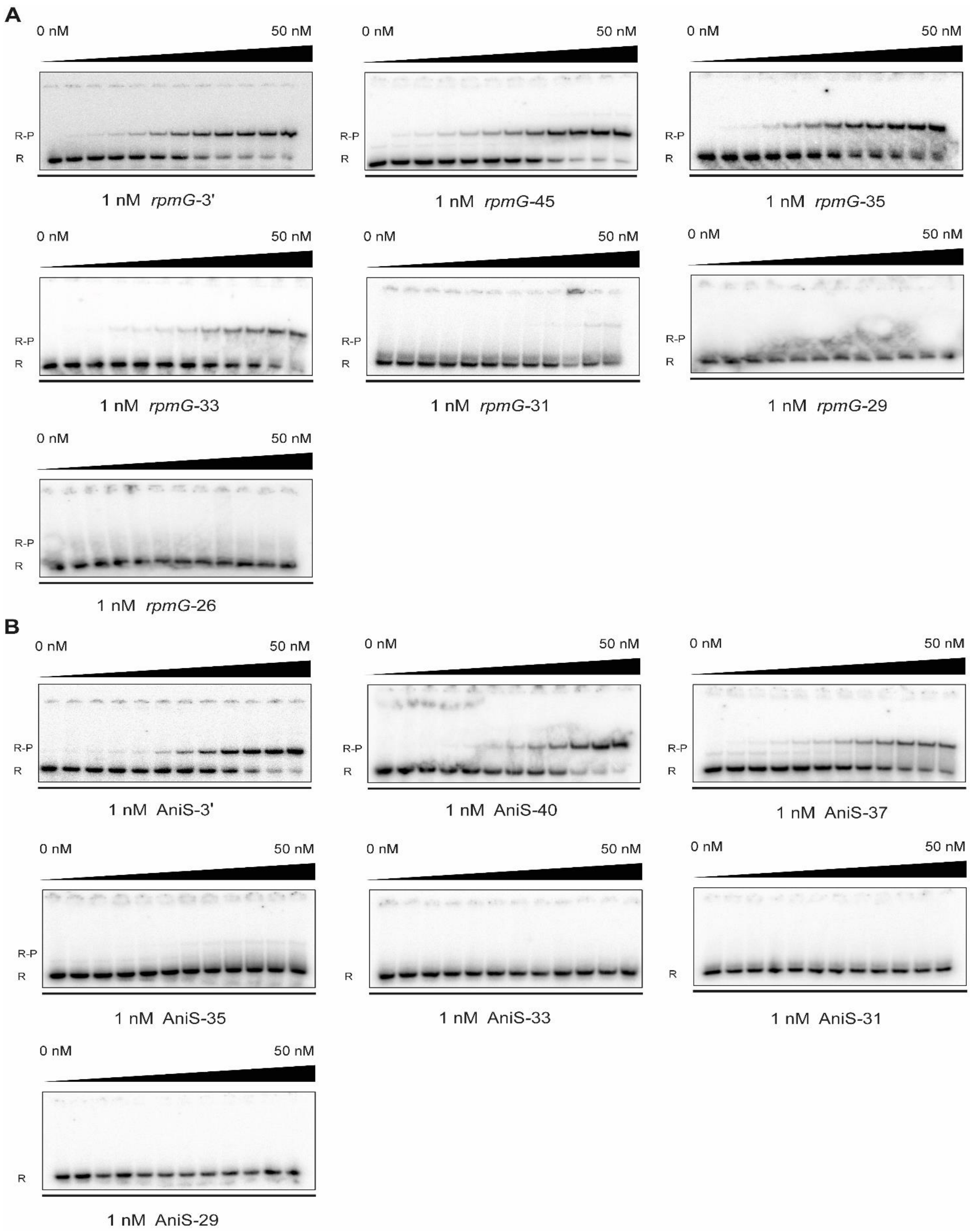


**Supplemental Figure S7. Gelshift analysis of the binding of $^{32}$P-labeled RNAs (A) *rpmG*-3', *rpmG*-45, *rpmG*-35, *rpmG*-33, *rpmG*-31, *rpmG*-29 and *rpmG*-26, (B) AniS-3', AniS-40, AniS-37, AniS-35, AniS-33, AniS-31 and AniS-29 to *N. meningitidis* ProQ.** The raw data in the gels correspond to the data presented in plots in Fig. 4 in the main text. The concentration series of ProQ protein was made by 2-fold sequential dilutions, and its range is indicated above the gels. Free $^{32}$P-RNA is marked as R, RNA-ProQ complex as R-P.

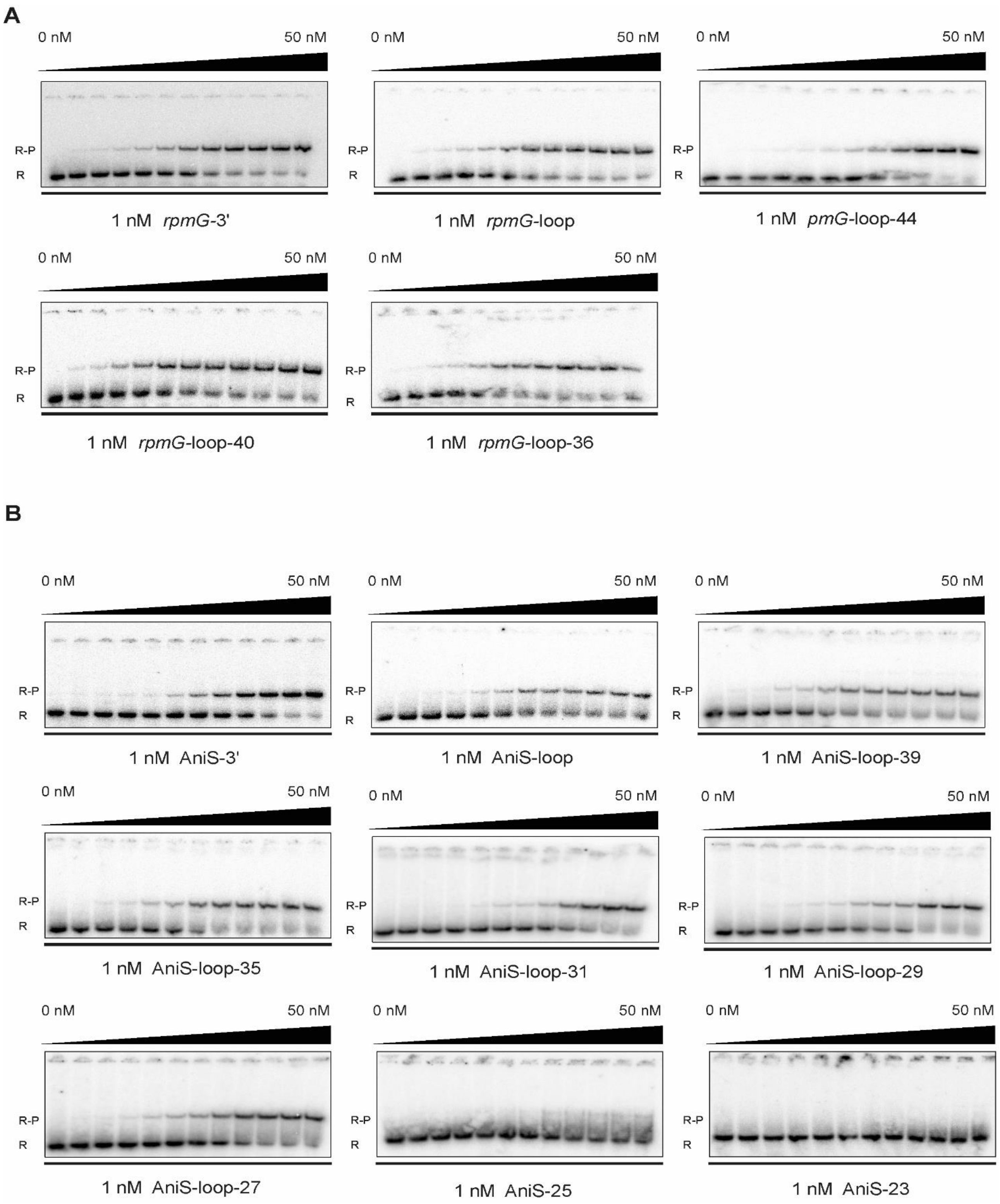


**Supplemental Figure S8. Gelshift analysis of the binding of $^{32}$P-labeled RNAs (A) *rpmG*-3', *rpmG*-loop, *rpmG*-loop-44, *rpmG*-loop-40 and *rpmG*-loop-36, (B) AniS-3', AniS-loop, AniS-loop-39, AniS-loop-35, AniS-loop-31, AniS-loop-29, AniS-loop-27, AniS-25 and AniS-23 to *N. meningitidis* ProQ.** The raw data in the gels correspond to the data presented in plots in Fig. 5 in the main text. The concentration series of ProQ protein was made by 2-fold sequential dilutions. The range of the ProQ concentrations is indicated above the gels. Free $^{32}$P-RNA is marked as R, RNA-ProQ complex as R-P.

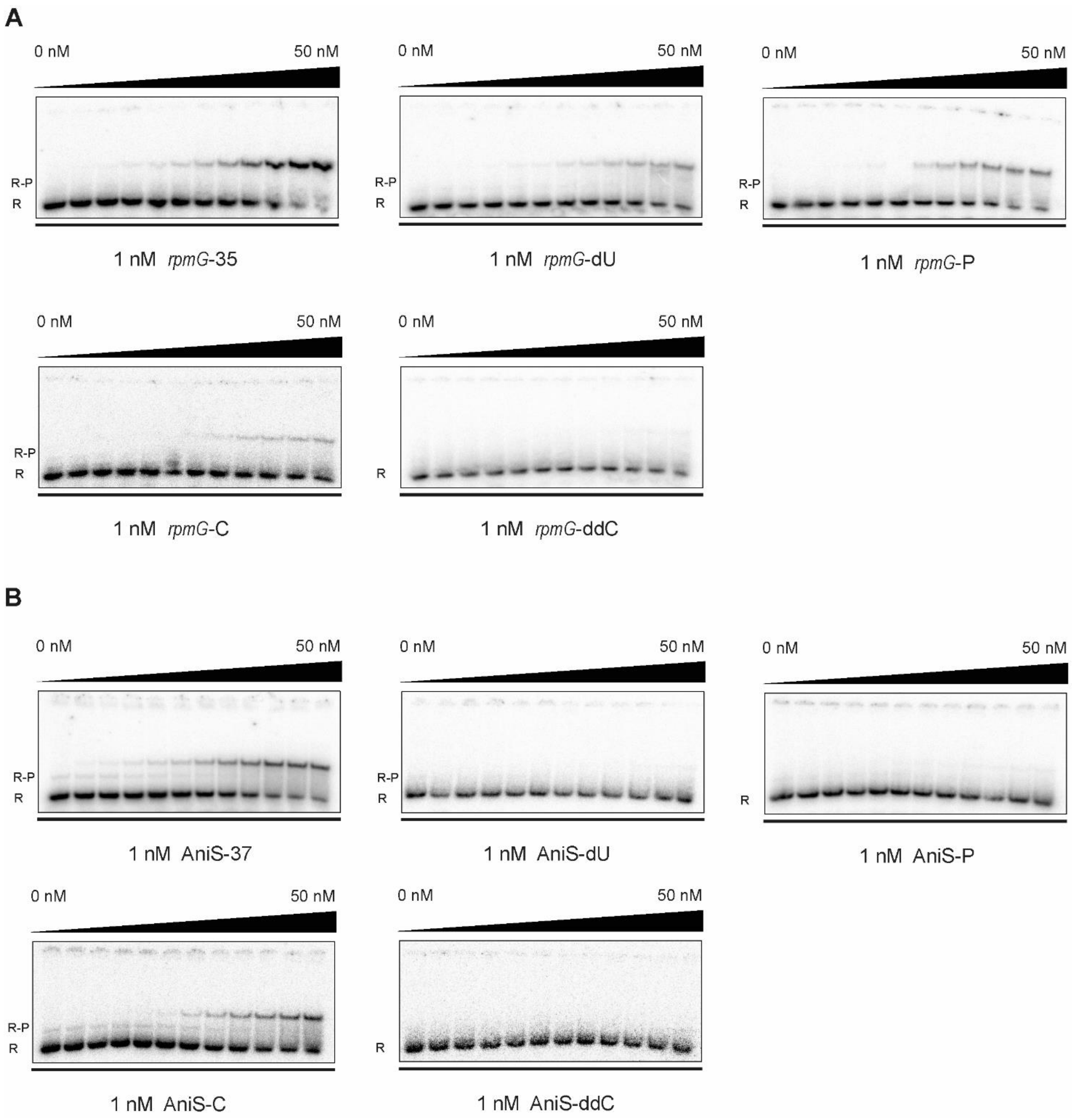


**Supplemental Figure S9. Gelshift analysis of the binding of $^{32}$P-labeled RNAs (A) *rpmG*-35, *rpmG*-dU, *rpmG*-P, *rpmG*-C and *rpmG*-ddC, (B) AniS-37, AniS-dU, AniS-P, AniS-C and AniS-ddC to *N. meningitidis* ProQ.** The raw data in the gels correspond to the data presented in plots in Fig. 6 in the main text. The concentration series of ProQ protein was made by 2-fold sequential dilutions. The range of the ProQ concentrations is indicated above the gels. Free $^{32}$P-RNA is marked as R, RNA-ProQ complex as R-P.

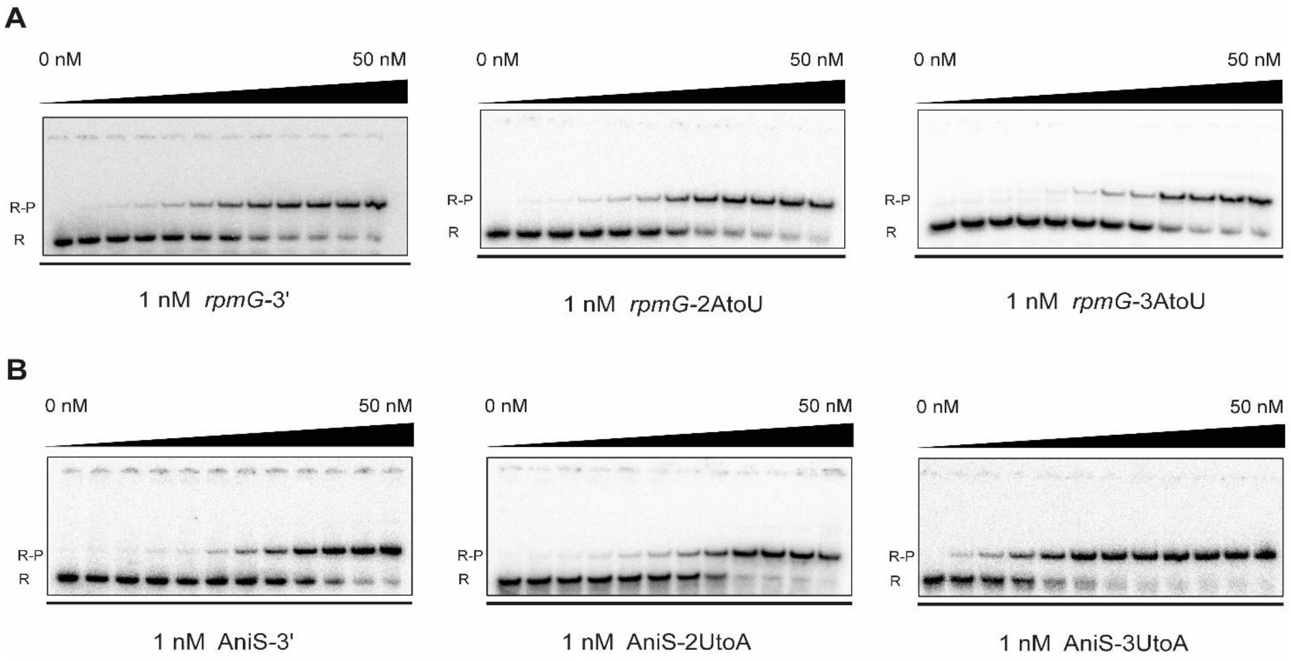


**Supplemental Figure S10. Gelshift analysis of the binding of $^{32}$P-labeled RNAs (A) *rpmG*-3', *rpmG*-2AtoU and *rpmG*-3AtoU, (B) AniS-3', AniS-2UtoA and AniS-3UtoA to the ProQ protein from *Neisseria meningitidis.*** The raw data in the gels correspond to the data presented in plots in Fig. 7 in the main text. The concentration series of ProQ protein was made by 2-fold sequential dilutions. The range of the ProQ concentrations is indicated above the gels. Free $^{32}$P-RNA is marked as R, RNA-ProQ complex as R-P.
